# Joint representation of spliced and unspliced RNA expands information content in single-cell expression data

**DOI:** 10.64898/2025.12.18.695314

**Authors:** Arya Jadhav, Zachary J. DeBruine

## Abstract

**Motivation:** Single-cell RNA-seq captures both mature (spliced) and nascent (unspliced) transcripts, yet standard preprocessing typically collapses these signals into a single expression matrix, obscuring recoverable transcriptional structure and introducing splicing-dependent artifacts in downstream embeddings in foundation models. We asked whether treating spliced and unspliced counts as distinct feature sets increases recoverable latent structure in unsupervised dimension reduction.

**Results:** Across four datasets with paired spliced and unspliced count layers, we applied non-negative matrix factorization (NMF) to spliced-only, unspliced-only, summed, and concatenated representations. Using cross-validation, we estimated the revealable rank *k*^∗^ (largest stable rank achieving near-optimal held-out error). In all datasets, concatenated representations consistently supported higher *k*^∗^ than spliced-only or unspliced-only inputs and typically exceeded the summed representation, indicating that preserving splicing status exposes additional stable latent programs. Minimal overlap between spliced- and unspliced-derived factors corresponded to modality-specific gene set enrichment patterns consistent with complementary steady-state and nascent regulatory activity. These results establish splicing status as an informative axis of variation and motivate preserving spliced and unspliced layers as first-class features to maximize information recovery in single-cell analysis.

**Availability and Implementation:** The `singlet` R package is available at github.com/zdebruine/singlet. Code to reproduce all figures is available at github.com/Arya-86/nmf-spliced-unspliced.

**Supplementary information:** Supplementary data are appended to this manuscript.

## Introduction

Single-cell RNA sequencing (scRNA-seq) has become a cornerstone technology for exploring cellular diversity and gene regulation at single-cell resolution. Its adoption has accelerated through large-scale initiatives such as the Human Cell Atlas [1] and the BRAIN Initiative Cell Census Consortium [2], which are generating reference atlases that will shape the next decade of transcriptomic discovery. As these datasets expand in scale and resolution, analytical conventions governing how gene expression is quantified and reduced have a profound influence on what biological structure can be revealed [3].

In most workflows, sequencing reads are aligned to a reference transcriptome to quantify spliced, mature mRNA molecules. The resulting gene expression matrix therefore reflects the abundance of processed transcripts that have completed splicing and export. However, single-cell sequencing protocols also capture a substantial proportion of unspliced, intron-containing pre-mRNAs [4]. These reads originate from nascent transcripts within the nucleus and carry information about genes currently undergoing transcriptional activation. Despite their ubiquity, unspliced reads are often discarded or summed with spliced reads, erasing distinctions between transcriptional activity and steady-state expression. This convention emerged largely from computational convenience and compatibility with downstream tools rather than empirical evidence of redundancy between the two signals [5].

The treatment of spliced and unspliced reads in scRNA-seq quantification has evolved markedly over the past decade. Early single-cell pipelines restricted quantification to reads mapping to exons, assuming that exon-aligned reads best represent functional mRNA abundance. With the advent of single-nucleus RNA-seq (snRNA-seq) and dynamic modeling frameworks, inclusion of intronic reads has become increasingly common. In such datasets, intronic reads can account for 20–50% of total unique molecular identifiers (UMIs) [6]. Modern quantification pipelines such as 10x Genomics `Cell Ranger` (v7.0 and later) and `STARsolo` now default to whole-transcriptome quantification, aligning reads to both exons and introns to maximize sensitivity and preserve information from both transcriptional compartments [7, 8]. This shift effectively redefines the standard measure of “gene expression” in scRNA-seq from cytoplasmic mRNA abundance to a composite signal reflecting total transcriptional output, even though intronic and exonic reads are typically aggregated.

The most influential framework exploiting splicing information is RNA velocity [4, 9]. RNA velocity estimates the future transcriptional state of a cell by comparing spliced and unspliced transcript abundances, modeling the kinetics of transcription, splicing, and degradation. Velocity analysis depends on constructing separate spliced and unspliced count matrices using specialized quantification tools such as `velocyto` [4], `alevin-fry` [10], or the velocity feature of `STARsolo` [8], which are now distributed as part of standard pipelines. While RNA velocity has produced compelling perspectives on cellular differentiation and lineage trajectories, it relies on simplifying assumptions regarding kinetic steady state, uniform splicing rates, and accurate classification of transcript status, which may introduce variability across datasets [11, 9]. Benchmark studies further demonstrate that velocity estimates can differ substantially depending on quantification software and parameterization [10, 12].

Nevertheless, the widespread adoption of RNA velocity has established a robust infrastructure for generating and storing spliced and unspliced matrices as standard outputs of single-cell pipelines. This availability enables direct multivariate analysis of these modalities as complementary inputs for exploratory analysis. Treating spliced and unspliced counts as distinct yet interrelated features may reveal additional structure in the data without invoking kinetic modeling.

The present study is motivated by a foundational question that remains unexplored despite the ubiquity of spliced and unspliced data: does treating spliced and unspliced transcripts as independent, first-class features enhance the latent structure recoverable from single-cell expression matrices? If these transcript classes encode complementary facets of gene regulation, then modeling both explicitly should increase the effective dimensionality and explainability of low-rank representations.

To investigate this question, we employ a cross-validated, unsupervised factorization framework based on Non-negative Matrix Factorization (NMF) [13, 14]. NMF provides an interpretable, additive decomposition well-suited to non-negative count-based biological data and allows direct evaluation of reconstruction fidelity and rank stability. By applying this framework to public scRNA-seq datasets quantified for both spliced and unspliced reads, we assess how these transcript classes jointly define the expressible subspace of transcriptional variation, extending splicing-aware analysis beyond trajectory inference toward a more general characterization of transcriptional structure.

## Results

### Revealable rank provides an interpretable measure of information content across feature representations

Single-cell RNA sequencing captures both mature (spliced) and nascent (unspliced) RNA molecules [4, 9]. Conventional workflows often collapse these signals into a single expression matrix, implicitly treating them as redundant. However, spliced and unspliced counts reflect different transcriptional processes and may therefore encode complementary biological information [11]. To test this hypothesis, we analyzed paired spliced and unspliced count layers across multiple datasets under four feature representations: spliced-only, unspliced-only, summed (spliced + unspliced), and concatenated (spliced ⊕ unspliced) features (Figure 1).

**Fig. 1:**
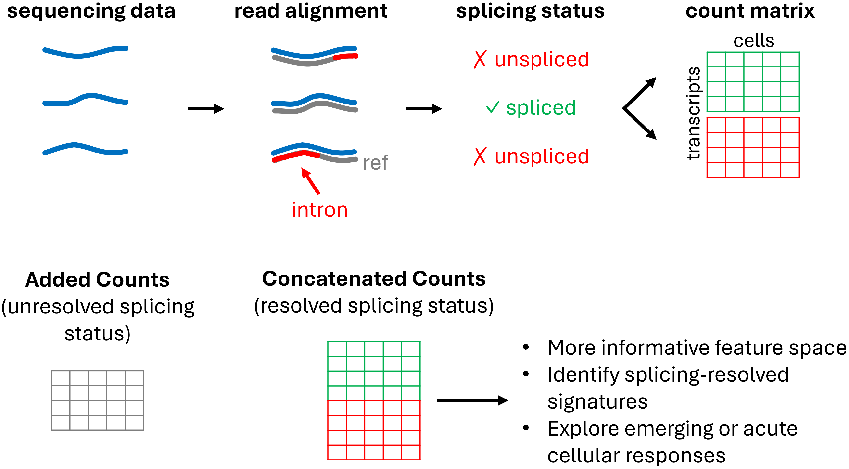
Conceptual overview of spliced and unspliced quantification in single-cell RNA-seq. Single-cell workflows typically align reads to transcriptomic references that do not include intronic regions, producing a count matrix of transcripts × cells that conflates spliced and unspliced molecules. When alignment includes intronic annotation, the splicing state of each read can be resolved, yielding two matrices—spliced and unspliced—that can be concatenated into a joint representation. This joint feature space provides a richer foundation for dimensionality reduction and integrative modeling of cellular state than standard pre-processing that adds spliced and unspliced reads together.

To quantify how feature representation affects the amount of structure recoverable from the data, we determine the revealable rank *k*^∗^: the largest factorization rank that can be learned stably without overfitting, as assessed by cross-validation [15, 16]. Intuitively, *k*^∗^ approximates the effective dimensionality of the transcriptional signal supported by a given representation, or in other words, the number of distinct latent programs that can be reconstructed in a way that generalizes beyond the observed entries.

We estimate *k*^∗^ by fitting NMF across a span of ranks and tracking held-out reconstruction error as model capacity increases. As illustrated in Figure 2B, low ranks underfit (high test set error that decreases with rank), intermediate ranks achieve stable minima, and sufficiently large ranks overfit (held-out error rises while training loss continues to decrease). We take *k*^∗^ as the largest rank prior to sustained overfitting, allowing that some intermediate ranks may be less stable than nearby higher ranks due to the metastability of high-dimensional transcriptional programs at increasing model capacity [17]. In the following section, we apply this framework uniformly to the four feature representations.

**Fig. 2:**
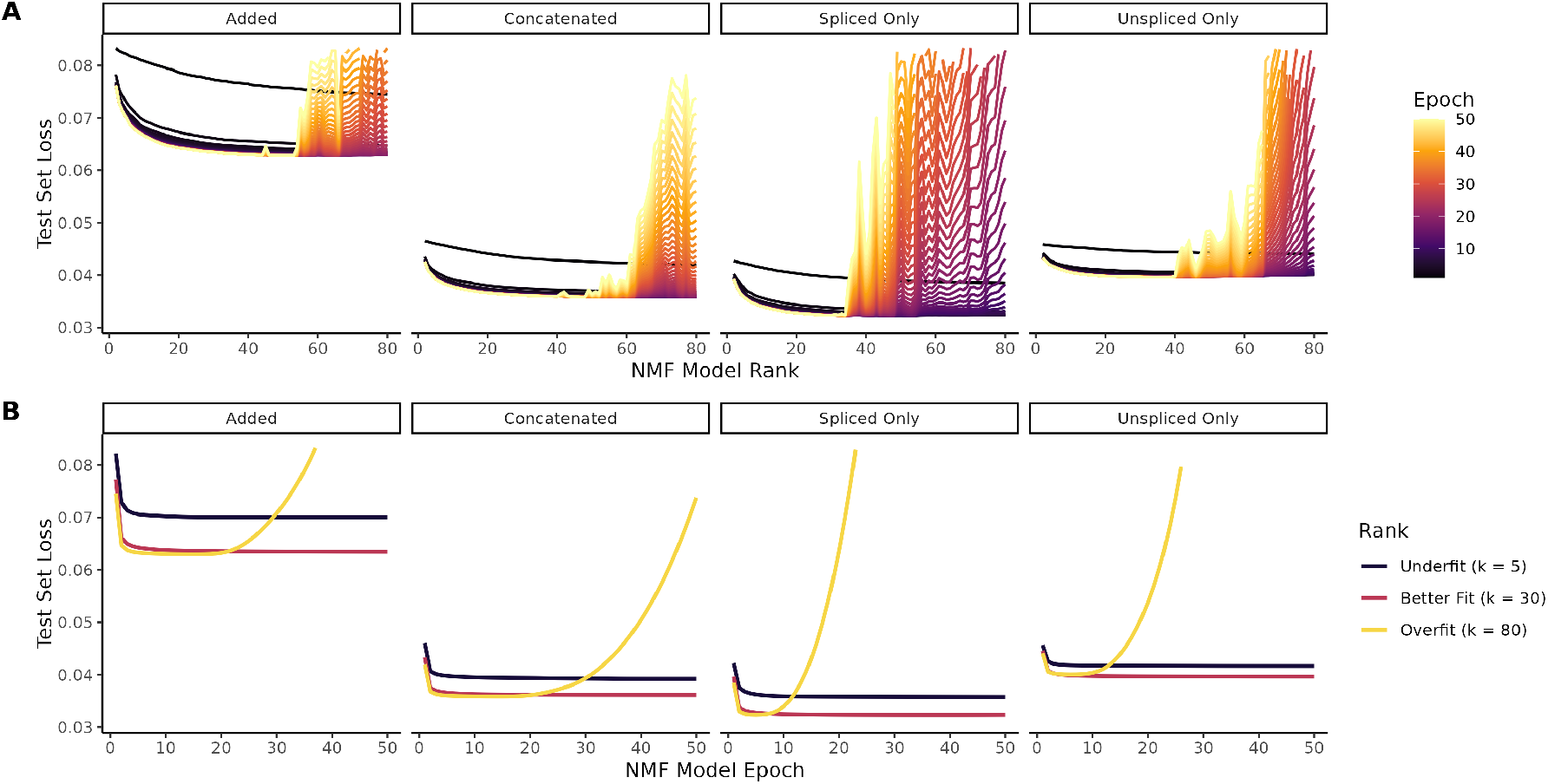
Cross-validation of NMF on the Heart dataset defines the revealable rank for each modality. (A) Mean test-set reconstruction error across NMF ranks (*k* = 2–80) for spliced-only, unspliced-only, added, and concatenated inputs. Low ranks underfit, showing large unexplained variance, whereas high ranks overfit, reflected by rising test error despite improved training loss. (B) Representative trajectories illustrate underfit (*k* = 10), approximate representative optimal (*k* = 30), and representative overfit (*k* = 80) behavior, demonstrating how overfitting is recognized as an instability of test loss as the model fits.

### Concatenating spliced and unspliced features increases revealable rank

We analyzed four public 10x Genomics datasets commonly used in RNA-velocity studies [11] (Heart, Brain, Neuron, and PBMC) to assess how spliced/unspliced feature representation affects the recoverable latent structure. Across all four datasets and three replicates, concatenating spliced and unspliced features generally increased the revealable rank relative to models fit to spliced-only or unspliced-only inputs (Figure 2A; Table 1; Figure S2; Table S1-4). The relative information content of the two modalities was dataset-dependent: in some datasets unspliced supported higher rank than spliced, whereas in others the reverse held, indicating that neither modality is uniformly dominant and both contain substantial resolvable information structure. Combining modalities increased revealable rank beyond either single layer but by less than the sum of both. Concatenation yielded equal or higher ranks than the added representation in all datasets (with PBMC differences within uncertainty), indicating that preserving spliced and unspliced counts as distinct features enables NMF to resolve additional latent programs that are partially obscured when the modalities are summed.

**Table 1.** Optimal NMF rank based on cross-validation results (mean ± SD across three replicates). Concatenation consistently increased revealable rank across all datasets, demonstrating that joint representation of spliced and unspliced information adds recoverable latent structure.

| Dataset | Spliced | Unspliced | Added | Concat |
| --- | --- | --- | --- | --- |
| Brain | 33.0 ± 1.0 | 39.7 ± 2.5 | 42.7 ± 2.5 | 46.3 ± 5.0 |
| Heart | 35.7 ± 2.9 | 49.7 ± 8.4 | 55.7 ± 3.8 | 56.3 ± 4.0 |
| Neuron | 63.0 ± 6.6 | 42.3 ± 3.1 | 70.7 ± 4.0 | 75.7 ± 3.2 |
| PBMC | 34.3 ± 6.0 | 25.7 ± 0.6 | 42.0 ± 2.6 | 38.0 ± 7.9 |

Principal Component Analysis (PCA) did not show a corresponding separation between spliced, unspliced, summed, and concatenated representations when assessed using scree plots of marginal explained variance (Figure S1). Unlike the cross-validated framework used to estimate revealable rank for NMF, PCA scree plots provide a descriptive heuristic and do not directly assess generalization or overfitting. More comparable evaluations would require cross-validated PCA procedures such as JackStraw resampling, which are computationally challenging at the scale of the datasets analyzed here. Accordingly, we restrict PCA comparisons to the scree plot and emphasize that the rank expansion observed for concatenated representations arises under an explicitly cross-validated factorization framework.

### Increased revealable rank reflects complementary latent structure between spliced and unspliced features

To determine whether the increase in revealable rank arises from genuinely complementary structure rather than redundant signal, we quantified the similarity between latent factors learned under each feature representation. We computed pairwise cosine similarity between NMF factors derived from spliced-only, unspliced-only, added, and concatenated inputs. Factors learned from spliced and unspliced data showed minimal overlap (Figure 3A, Figure S3A), indicating that the two modalities capture largely distinct transcriptional programs.

**Fig. 3:**
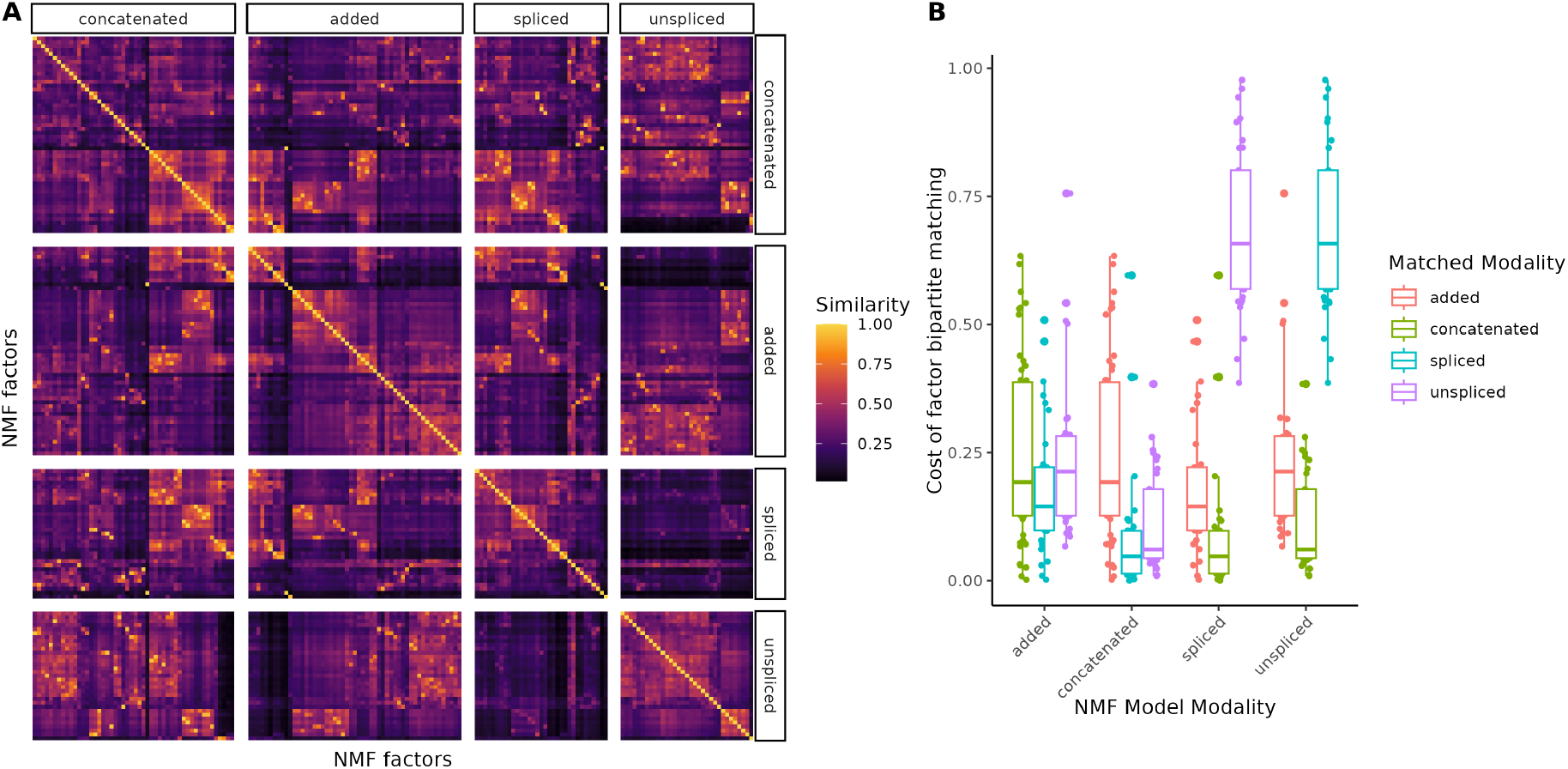
Spliced and unspliced modalities encode complementary latent structure. (A) Pairwise cosine similarity between NMF factors learned from spliced-only, unspliced-only, added, and concatenated inputs. Factors derived from spliced and unspliced data from the heart dataset exhibit minimal overlap, indicating largely non-redundant structure. (B) Optimal bipartite matching of factors across models from the heart dataset based on the similarity matrix in (A). Factors from the concatenated model show stronger correspondence to both spliced- and unspliced-derived factors than those from the added model, demonstrating that concatenation preserves modality-specific latent programs that are partially obscured by summation.

Both added and concatenated models exhibited partial correspondence to factors learned from the individual modalities. However, optimal bipartite matching revealed a clear distinction between these representations: factors from the concatenated model matched more closely to both spliced- and unspliced-derived factors than those from the added model (Figure 3B; Figure S3B). This indicates that summation partially entangles modality-specific signals, whereas concatenation preserves them as separable latent dimensions. These findings are likewise reflected on UMAP embeddings of the four modalities where concatenation yields the best-resolved substructure (Figure S4). Together, these results explain why concatenating spliced and unspliced features supports higher revealable rank—by retaining complementary structure that is obscured when the two modalities are collapsed.

### Modality-specific NMF factors correspond to distinct biological programs

To assess whether the structural separation between spliced and unspliced factors corresponds to meaningful biological differences, we examined the relative contribution of each modality to individual latent factors and evaluated their functional enrichment. For each factor in the concatenated model, we quantified the proportion of total feature loadings attributable to spliced versus unspliced transcripts and performed Gene Set Enrichment Analysis (GSEA) independently on each modality.

We used MSigDB Gene Ontology (C5) terms for GSEA, subsetting MSigDB pathways to the gene universe present in the single-cell transcriptomics data, and vice versa. Many latent factors had a preference for one modality over the other, indicating that NMF assigns distinct transcriptional signals to separable dimensions (Figure 4A).

**Fig. 4:**
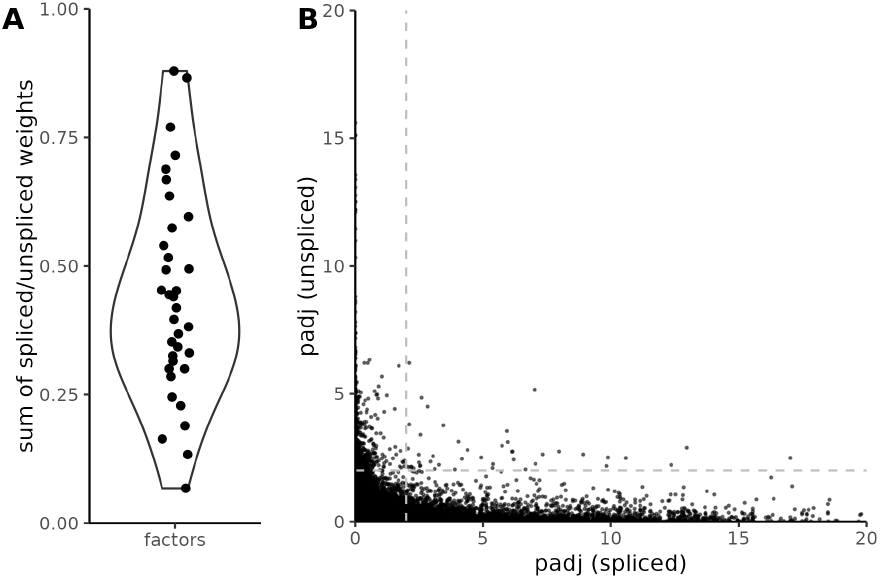
NMF factors show modality preference and learn distinct biological programs between spliced and unspliced features. (A) Proportion of spliced versus unspliced feature loadings per latent factor in the optimal-rank NMF model of the heart dataset, showing that many factors have a modality preference. (B) BH–adjusted p-values from GO C5 Gene Set Enrichment Analysis (GSEA) performed independently on spliced and unspliced loadings within each factor of the heart dataset shows no correlation, indicating that spliced and unspliced features capture biologically distinct transcriptional programs even within factors.

Consistent with this factor-specific modality preference, biological enrichment patterns differed markedly between spliced and unspliced features within the same factor. Across factors, we observed no correlation between BH–adjusted GSEA p-values derived separately from spliced versus unspliced gene weights (Figure 4B), demonstrating that the two modalities emphasize distinct regulatory programs rather than redundant biological processes. Together, these results show that preserving spliced and unspliced features yields latent factors that are not only structurally distinct but also biologically interpretable, providing a functional explanation for the increased revealable rank observed when the modalities are modeled jointly.

### Concatenated latent factors exhibit regulatory complementarity between spliced and unspliced signals

In addition to factors dominated by a single modality, the concatenated model learned latent dimensions in which spliced and unspliced features contributed substantially within the same factor. To assess whether these shared factors reflected coherent biological programs, we performed GSEA separately on spliced and unspliced loadings for each factor. Across multiple factors, spliced and unspliced features within the same latent dimension highlighted distinct biological processes, indicating complementary regulatory signatures captured within a single factor (Figure 5).

**Fig. 5:**
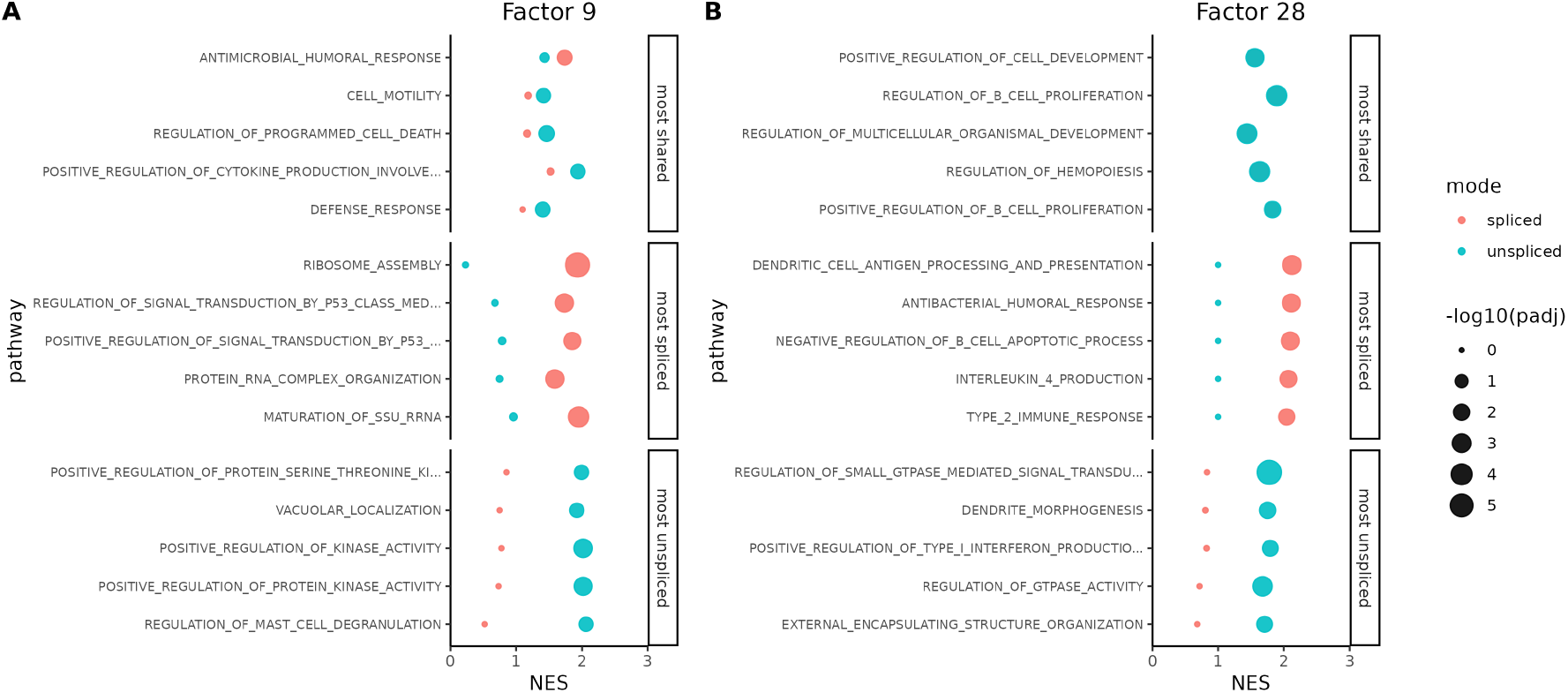
Concatenated latent factors encode complementary regulatory signatures across spliced and unspliced signals. Gene Set Enrichment Analysis (GSEA) performed separately on spliced and unspliced feature loadings within selected latent factors from the concatenated model. (A) Factor 9 from the heart dataset shows unspliced enrichment for signaling and mast-cell–associated pathways, whereas spliced enrichment emphasizes ribosome assembly and p53-related processes. (B) Factor 28 exhibits a similar separation, with unspliced features enriched for morphogenesis- and future-state programs and spliced features reflecting immune activation.

For example, in Factor 9 of the heart dataset, unspliced loadings were enriched for kinase signaling and mast-cell degranulation, whereas spliced loadings emphasized ribosome assembly and p53 signaling. Similarly, in Factor 28, unspliced enrichments captured programs associated with morphogenesis and cell-state specification, while spliced enrichments reflected immune activation and translational activity. These patterns generalized across factors (more examples presented in Figure S5), with unspliced features frequently enriched for transcriptional activation, signaling, and developmental pathways, whereas spliced features preferentially captured translation, oxidative metabolism, cell-cycle regulation, and other homeostatic processes.

Together, these results show that concatenating spliced and unspliced transcripts enables a single latent factor to encode multiple, functionally distinct aspects of a biological program, without requiring explicit kinetic or temporal modeling. By preserving both modalities, NMF captures complementary regulatory signatures that are otherwise conflated when transcript origin is ignored.

## Discussion

A central result of this study is that preserving splicing status in single-cell RNA-seq data materially increases the amount of latent structure that can be stably recovered. By treating spliced and unspliced transcripts as independent features, rather than collapsing them into a single expression matrix, we show that common single-cell datasets support a slightly higher revealable rank under cross-validated NMF. This increase reflects genuine, explainable structure rather than overfitting, indicating that standard preprocessing conventions discard resolvable information before analysis begins.

Our findings clarify that spliced and unspliced counts encode complementary, not redundant, signals. Across datasets, factors learned from spliced and unspliced layers exhibited minimal overlap, while concatenated models preserved modality-specific structure more faithfully than additive representations. These results establish that summation obscures distinctions that low-rank models can otherwise resolve when modalities are presented jointly. From a representation perspective, splicing status functions as an informative axis of variation rather than a nuisance variable.

Importantly, the additional structure exposed by concatenation is biologically interpretable. Factors dominated by spliced or unspliced features preferentially enriched for distinct gene programs, and even within shared factors, the two modalities highlighted different aspects of the same biological processes. These patterns are consistent with unspliced transcripts reflecting transcriptional activation and nascent responses, while spliced transcripts emphasize established or steady-state programs. Without invoking kinetic assumptions, this dual representation provides a richer and more nuanced view of cellular state.

Methodologically, we introduce revealable rank as a practical, cross-validated measure of information content that enables principled comparison across feature representations. While our analysis focuses on NMF, the core insight is representation-centric rather than model-specific: preserving splicing status expands the expressible subspace available to downstream models.

These findings carry immediate implications for large-scale single-cell atlases. As community resources increasingly standardize preprocessing and archive expression matrices for reuse, decisions about whether to collapse or preserve splicing layers directly affect the biological structure that future analyses can uncover. Retaining spliced and unspliced counts as first-class features substantially increases analytical flexibility, interpretability, and downstream reuse potential.

In summary, splicing status is not merely auxiliary metadata for dynamic modeling, it is a foundational component of transcriptional signal. Preserving spliced and unspliced layers expands the dimensionality of discoverable biology without altering experimental design or requiring new assays. For atlas builders, method developers, and curators of public single-cell resources, this work argues for a simple but consequential shift in practice: store more of what the data already contain, and let downstream models decide what structure can be revealed.

## Methods

### Single-cell RNA-seq datasets

We analyzed four 10x Genomics scRNA-seq datasets (PBMC, heart, brain, neuron) considered in Gorin *et al*. [11]. In their study, datasets were obtained from 10x Genomics and processed with standard preprocessing workflows to quantify spliced and unspliced molecules. Gorin *et al*. made available scripts and loom files generated by these workflows, which we used. In this work, we did not reprocess raw sequencing data or regenerate spliced/unspliced counts; instead, we used the released count matrices exactly as provided. Loom containers were parsed in R via `reticulate`, and spliced and unspliced layers were exported to `.npz` format before being cast to sparse `dgCMatrix` objects using the `Matrix` package. Cells were retained if they contained between 200 and 15,000 detected features and between 500 and 75,000 total UMIs. Genes detected in fewer than 10 cells were removed. No additional feature selection or variance filtering was applied.

### Construction of spliced and unspliced feature matrices

From the raw spliced and unspliced count matrices, we constructed four modality-specific input matrices:

1. Spliced-only (*s*).
2. Unspliced-only (*u*).
3. Added (*s*+*u*): per-gene alignment by gene symbol, followed by element-wise summation of the raw counts.
4. Concatenated (*s* ⊕ *u*): row-wise concatenation of the raw spliced and unspliced counts, preserving gene identity by appending modality-specific suffixes (`_s, _us`).

Gene sets were identical between spliced and unspliced matrices. No reweighting was applied between modalities, and no group-wise or batch normalization was performed. All four representations were analyzed in parallel in downstream experiments.

### Preprocessing and normalization

All input matrices were normalized using Seurat standard log-normalization with default parameters. No additional feature selection or variance filtering was applied for NMF. For PCA analyses, 2000 highly variable genes were selected and gene-wise centering and unit-variance scaling was applied using the standard Seurat workflow.

### Non-negative matrix factorization

For each dataset and each modality construction (*s, u, s*+*u, s* ⊕ *u*), we applied Non-negative Matrix Factorization (NMF) to decompose the normalized expression matrix 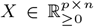 into a low-rank product

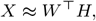

where *p* denotes the number of features, *n* the number of cells, and *k* the factorization rank. The matrix 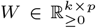 contains non-negative gene loadings for each factor, and 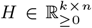 contains the corresponding non-negative cell loadings. We evaluated ranks *k* ∈ {2, 3, …, 80} for all datasets and modalities.

Models were fit by minimizing squared reconstruction error under non-negativity constraints,

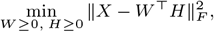

using an alternating least squares scheme with non-negative least squares updates. All factorization was performed using our open-source `singlet` implementation (C++ with R bindings), which is optimized for sparse inputs and parallel execution.

For each replicate, factor matrices were initialized from a uniform distribution on (0, 1) and trained for a fixed number of epochs (50), independent of dataset, modality, or rank. No regularizations, modality reweighting, or other stopping criteria were applied.

### Rank selection via cross-validation

To assess generalization performance and determine the effective dimensionality supported by each data representation, we employed an entry-wise cross-validation strategy based on sparse holdout masking, as implemented in the `cross_validate nmf` function in the `singlet` R package. For each dataset and modality, a fixed subset of matrix entries was designated as a test set. Each entry (*i, j*) was assigned to the held-out set with probability approximately 0.05, using a deterministic two-dimensional congruential pseudorandom number generator indexed by (*i, j*) and a replicate-specific seed. Masking was applied uniformly over all entries, including both zero and nonzero values.

Let Ω denote the set of held-out indices and Ω^*c*^ its complement. During model fitting, NMF updates were performed using only entries in Ω^*c*^, while performance was evaluated on Ω. The same mask was reused across all epochs for a given rank, and for all ranks in each replicate, ensuring that cross-validation comparisons reflected model capacity across ranks rather than stochastic resampling effects.

Model performance was quantified using mean squared reconstruction error on the held-out entries,

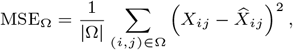

where 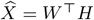 denotes the reconstructed matrix.

For each dataset, modality, and rank, we fit three independent replicates distinguished by random seeds. The same set of seeds was reused across ranks to ensure stable cross-rank comparisons. Cross-validation curves and downstream summaries report the mean held-out error across replicates.

### Determination of revealable rank

For each dataset, representation, and replicate, we fit NMF across a contiguous span of ranks and tracked the held-out reconstruction error MSE_Ω_(*k*) after convergence. For that replicate, we selected the revealable rank *k*^∗^ as the largest rank whose converged held-out error was near, or indistinguishable from, the minimum held-out error achieved over all tested ranks, and that lay immediately before the onset of pronounced overfitting at immediately greater ranks. Operationally, overfitting was recognized as a sharp and sustained rise in MSE_Ω_(*k*) with increasing *k* despite continued improvement of the training objective, producing a generally very sharp inflection point separating a broad near-minimum region from an overfit region. This selection was performed on the cross-validation curves for each replicate and dataset and is shown in Tables S1-S4. Reported *k*^∗^ values in the main text (Table 1) are the mean ± standard deviation of the replicate-specific selections (three replicates per condition).

### Factor similarity and cross-modality matching

To quantify correspondence between latent factors learned from different input modalities, we compared NMF gene-loading matrices across models. For each modality-specific model, we formed a factor-by-factor similarity matrix *S* between columns of the two gene-loading matrices (e.g., using dot products of *ℓ*_2_-normalized loading vectors, which is equivalent to cosine similarity). We then computed an optimal one-to-one correspondence between factors by solving a bipartite matching problem using `RcppML::bipartiteMatch`. Specifically, factors were matched to maximize total similarity (or equivalently to minimize a cost matrix derived from similarity, e.g., *C* = 1−*S*). All matching was performed on factors learned at the revealable rank for each dataset and modality.

### Gene set enrichment analysis

To interpret the biological programs captured by individual NMF factors, we performed Gene Set Enrichment Analysis (GSEA) on factor-specific gene loadings. For each factor, genes were ranked by their normalized loading weights in the gene-loading matrix *W*. In the concatenated modality, spliced and unspliced features were analyzed separately: GSEA was performed independently on the spliced-derived gene weights and the unspliced-derived gene weights associated with the same latent factor. Enrichment analysis was conducted using the `fgsea` package in R, with gene sets drawn from the MSigDB Gene Ontology C5 collection. The gene universe for each analysis was restricted to genes present in the corresponding spliced or unspliced layer after quality control. This strategy enables direct comparison of biological processes emphasized by spliced versus unspliced contributions within shared latent factors, facilitating interpretation of modality-specific and temporally complementary regulatory programs.

### Software and data availability

All datasets analyzed in this study are publicly available from the Caltech Data repository[18]. DOI: `10.22002/D1.20030`. All NMF analyses were performed using the `singlet` library, which provides a C++ implementation of non-negative matrix factorization with an R interface. The source code for `singlet` is publicly available at https://github.com/zdebruine/singlet, including the `cross_validate_nmf` function used in this study. Complete scripts required to reproduce all figures, cross-validation analyses, and downstream results presented in this manuscript are available at https://github.com/Arya-86/nmf-spliced-unspliced. All analyses were executed on CPU-only systems with multi-threaded execution (up to 128 threads); results were invariant to thread count in our implementation and do not necessarily require HPC resources due to the memory and computational efficiency of the `singlet` C++ implementation and relatively small size of these datasets. All random number generation was controlled via fixed seeds, and identical seeds were reused across ranks within each replicate to ensure deterministic and comparable cross-validation results. Given identical inputs, seeds, and software versions, all reported results are fully reproducible.

## Author contributions statement

Z.D. conceived the study, proposed the central hypothesis, designed the analytical framework, developed the `singlet` software library and NMF cross-validation methodology used in this work, contributed to figure generation, and wrote the manuscript. A.J. performed the data analysis, generated figures, and assembled the code used to produce the manuscript results and visualizations. Both authors contributed to the interpretation of results and approved the final manuscript.

## Acknowledgments

We thank the Chan Zuckerberg Initiative Single-cell Data Insights working group members for productive discussions on this issue and interest in understanding the value of providing both spliced and unspliced read counts in the Cell Census. We also refer to the GitHub issue `/chanzuckerberg/cellxgene-census/issues/291` which led to deeper attention to this topic.

