## Supplemental Material for "Joint representation of spliced and unspliced RNA expands information content in single-cell expression data"

### Supplementary Figure 1: PCA Scree Plots by Modality

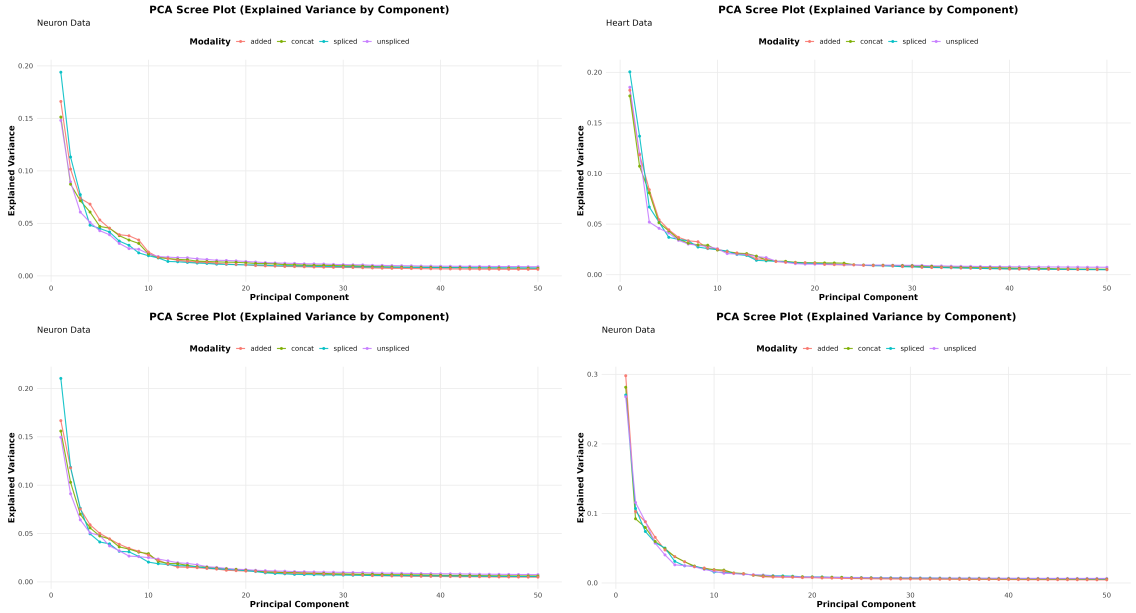


PCA scree plots (explained variance by component) for each dataset: Brain, Heart, Neuron, and PBMC. Modalities include spliced, unspliced, added, and concatenated. Note that differences in explained variance across modalities are minimal, suggesting PCA is relatively insensitive to transcript origin.

### Supplementary Figure 2: Cross-validation Results for other datasets

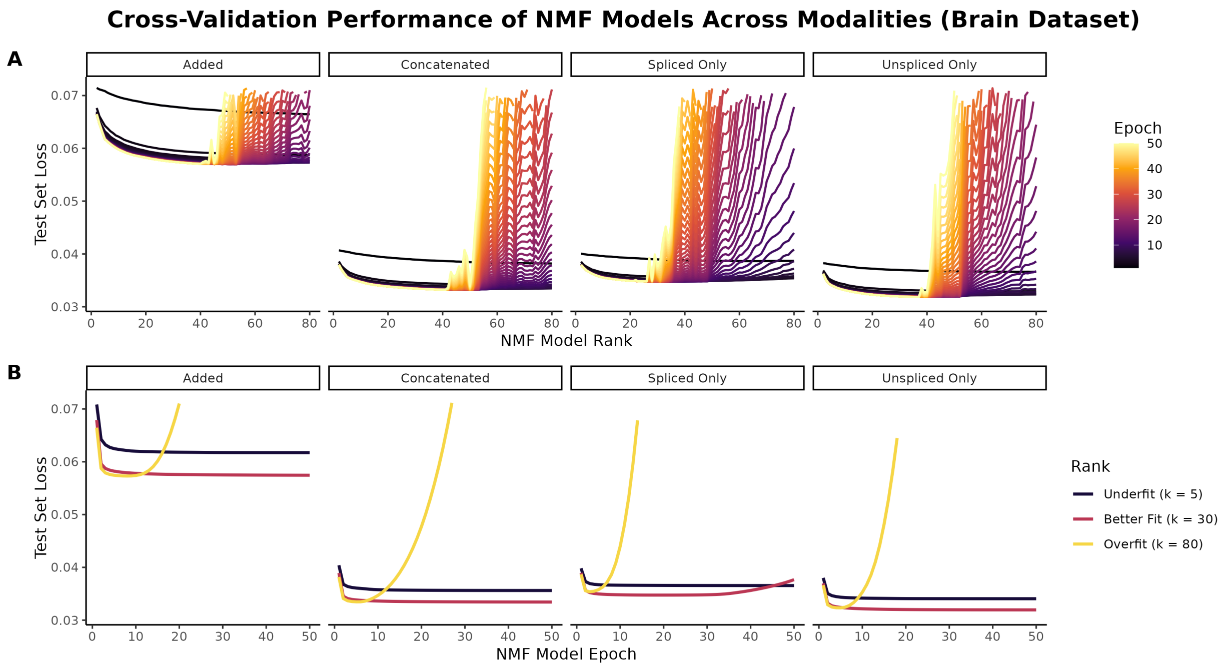


Brain dataset cross-validation results, as done in Figure 2 for the heart dataset.

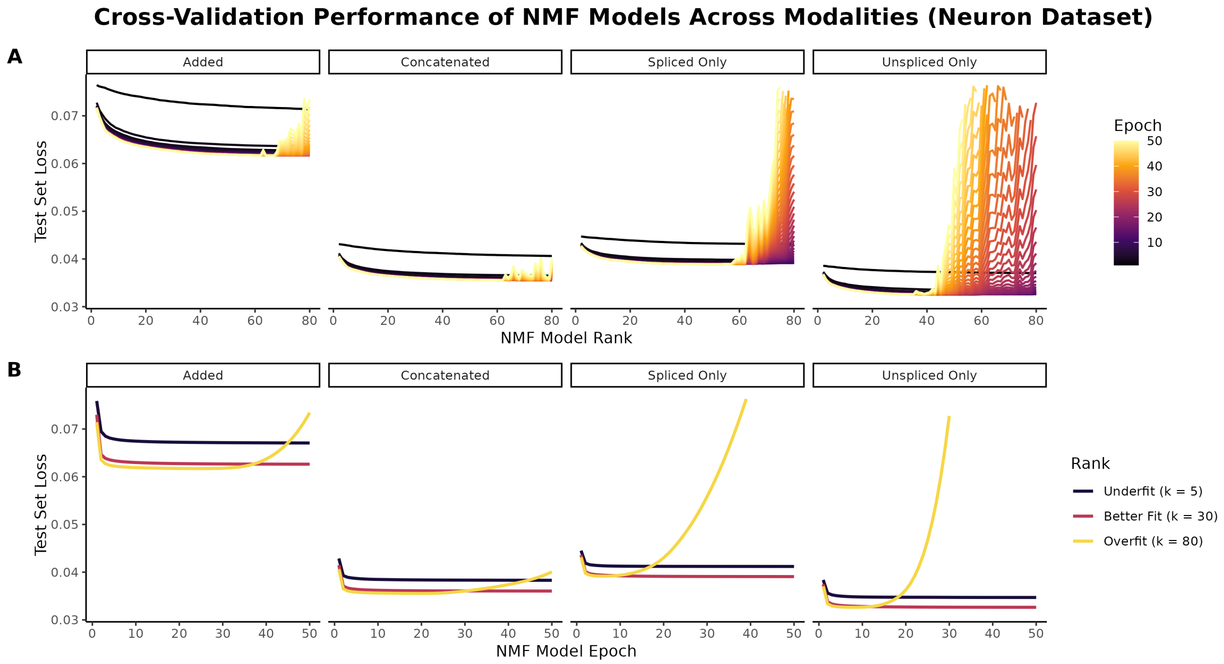


Neuron dataset cross-validation results, as done in Figure 2 for the heart dataset.

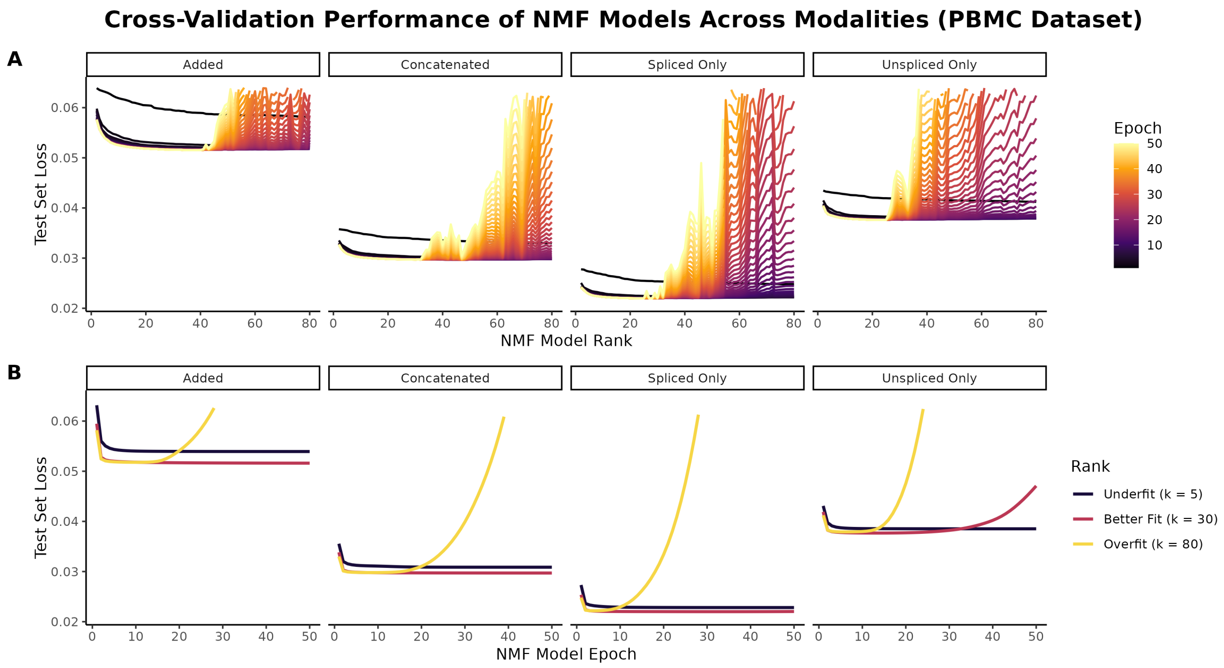


PBMC dataset cross-validation results, as done in Figure 2 for the heart dataset.

### Supplementary Figure 3: Factor matching results for other datasets

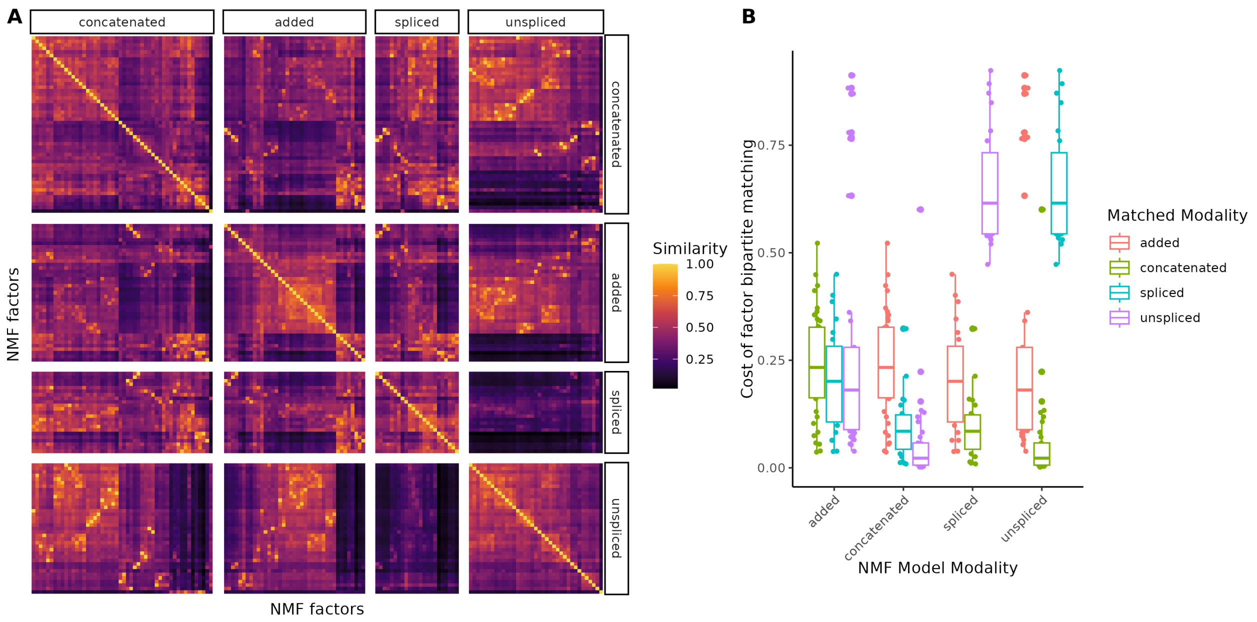


Brain dataset factor matching results, as done in Figure 3 for the heart dataset.

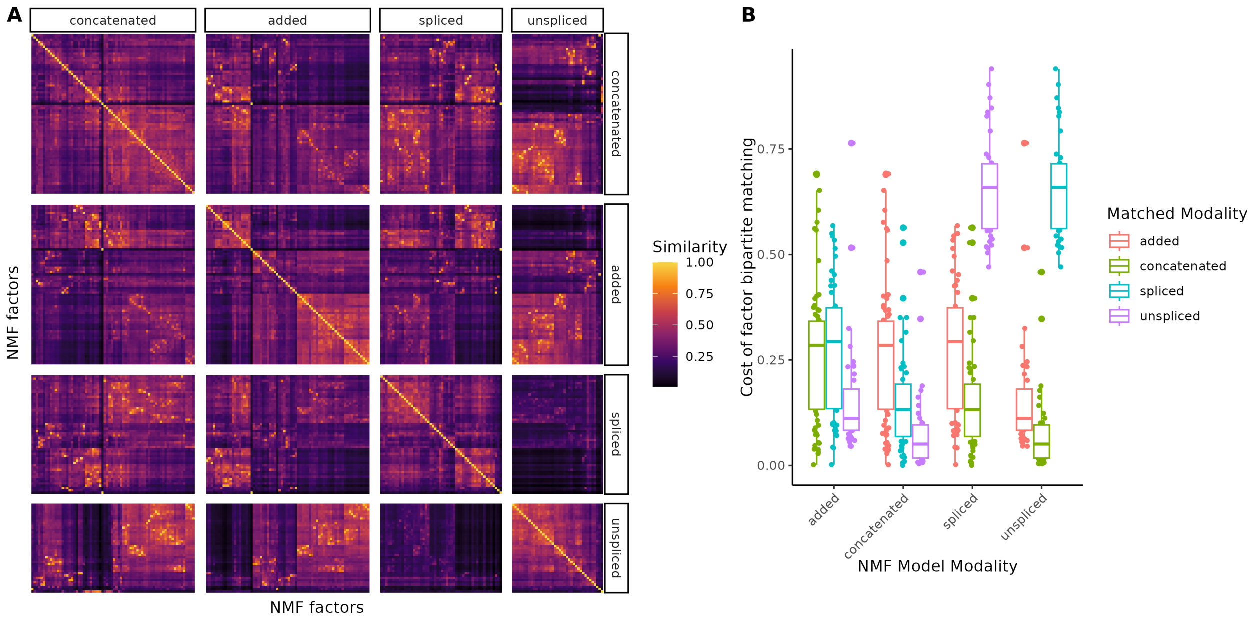


Neuron dataset factor matching results, as done in Figure 3 for the heart dataset.

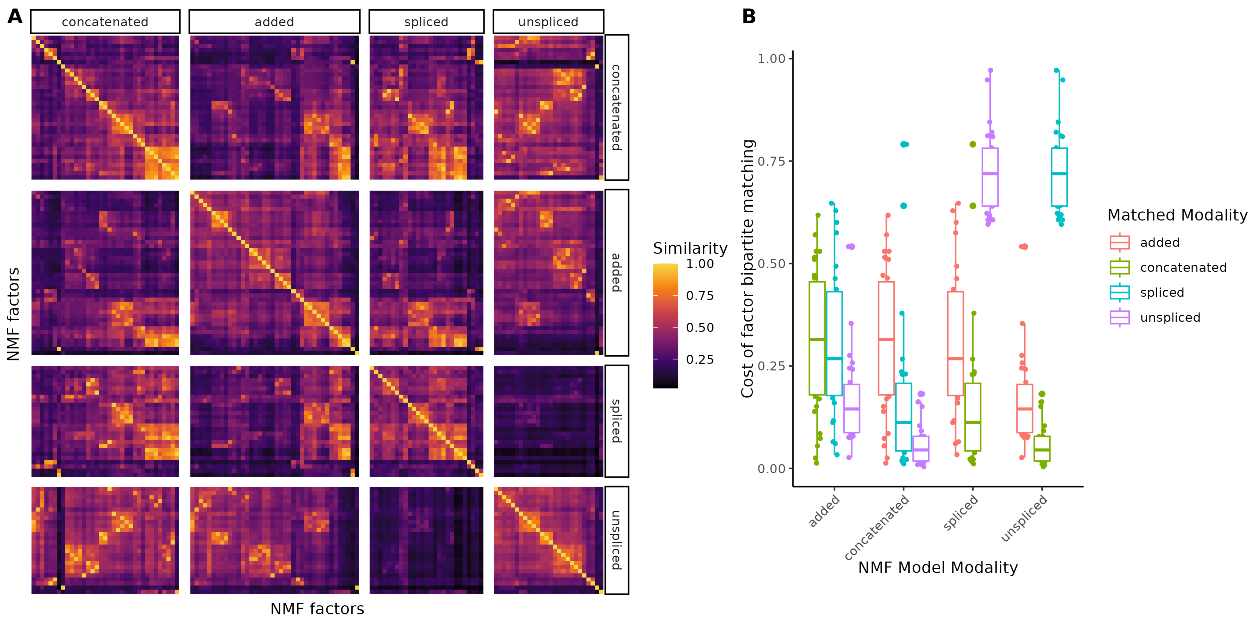


PBMC dataset factor matching results, as done in Figure 3 for the heart dataset.

#### Supplementary Figure 4: Comparison of PCA and NMF embeddings in UMAP space for heart dataset

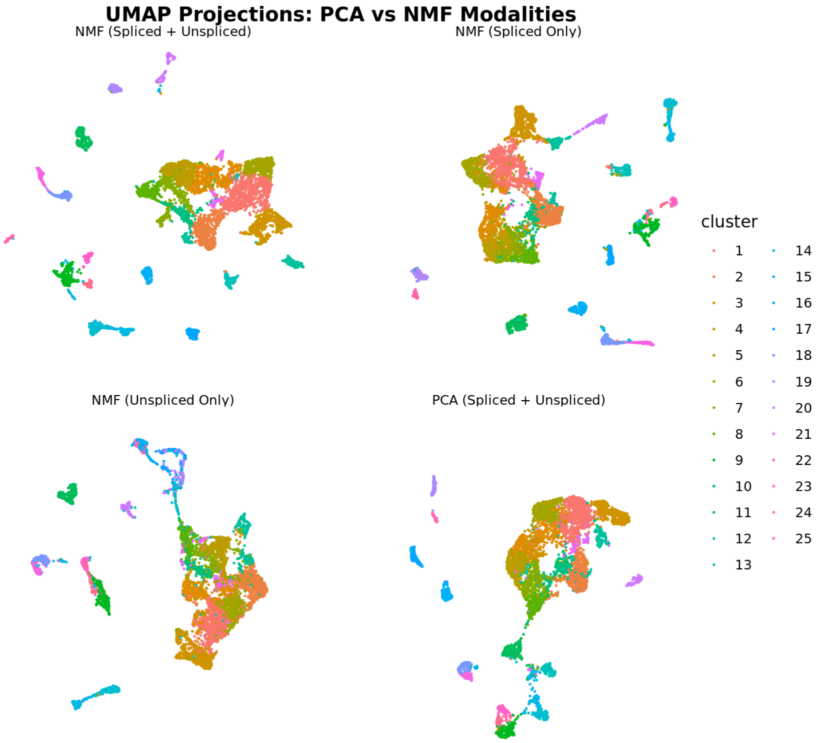


Comparison of UMAP embeddings for PCA and NMF models at optimal rank (PCA rank determined by inflection point in scree plot) showing that NMF models learn sharper embeddings regardless of what modality is used, likely attributable to higher revealable rank in NMF.

#### Supplementary Figure 5: GSEA Plots for Selected Additional PBMC Dataset NMF Factors (as done in Figure 5)

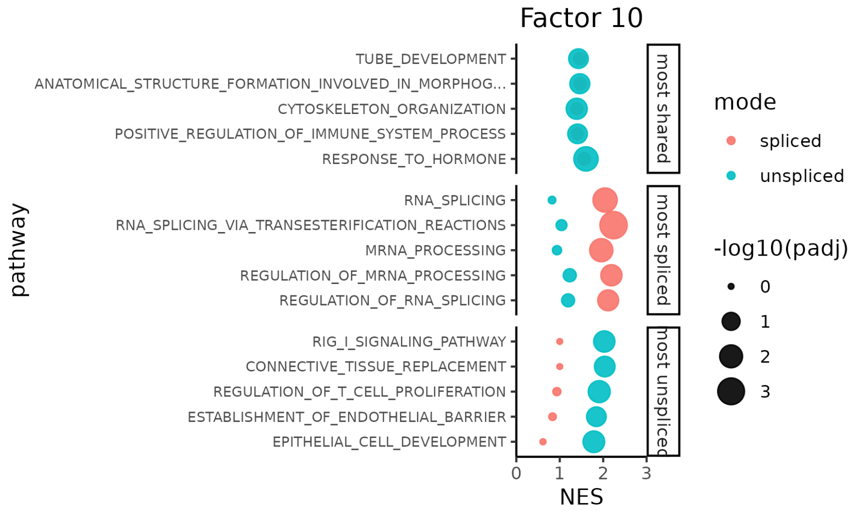


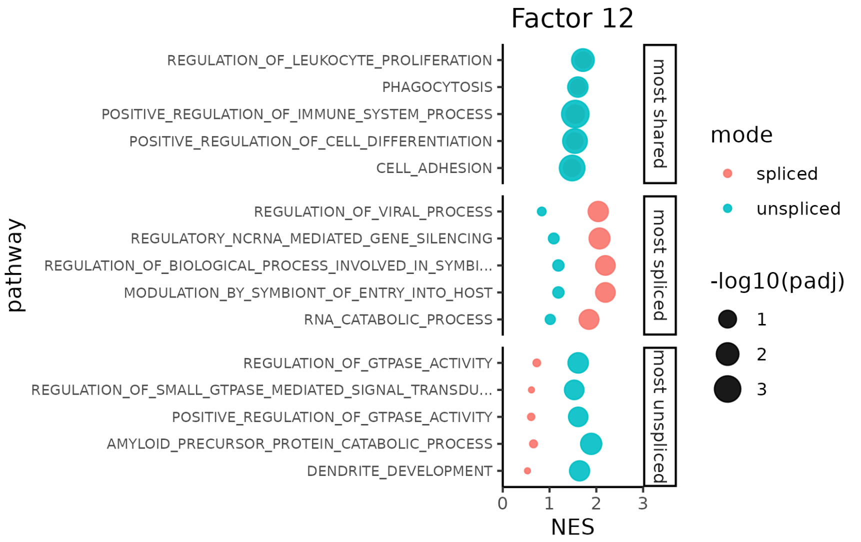

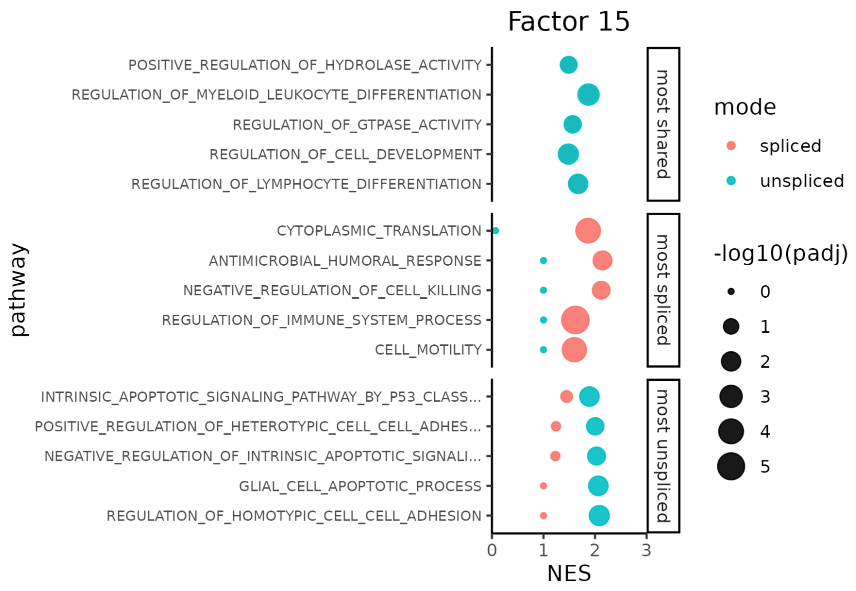
Factor 15
POSITIVE_REGULATION_OF_HYDROLASE_ACTIVITY
REGULATION_OF_MYELOID_LEUKOCYTE_DIFFERENTIATION
REGULATION_OF_GTPASE_ACTIVITY
REGULATION_OF_CELL_DEVELOPMENT
REGULATION_OF_LYMPHOCYTE_DIFFERENTIATION
CYTOPLASMIC_TRANSLATION
ANTIMICROBIAL_HUMORAL_RESPONSE
NEGATIVE_REGULATION_OF_CELL_KILLING
REGULATION_OF_IMMUNE_SYSTEM_PROCESS
CELL_MOTILITY
INTRINSIC_APOPTOTIC_SIGNALING_PATHWAY_BY_P53_CLASS...
POSITIVE_REGULATION_OF_HETEROTYPIC_CELL_CELL_ADHES...
NEGATIVE_REGULATION_OF_INTRINSIC_APOPTOTIC_SIGNALI...
GLIAL_CELL_APOPTOTIC_PROCESS
REGULATION_OF_HOMOTYPIC_CELL_CELL_ADHESION
most shared
most spliced
most unspliced
pathway
NES
0 1 2 3
mode
spliced
unspliced
-log10(padj)
0
1
2
3
4
5

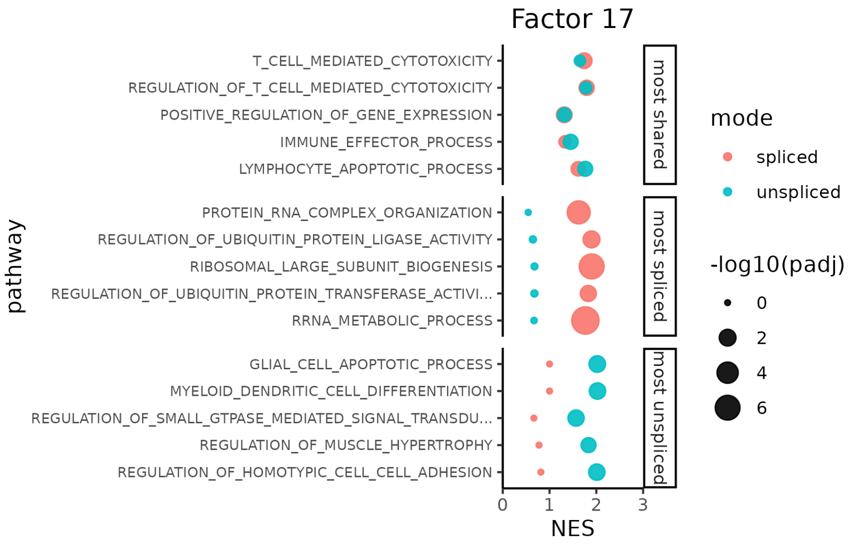
Factor 17
T_CELL_MEDIATED_CYTOTOXICITY
REGULATION_OF_T_CELL_MEDIATED_CYTOTOXICITY
POSITIVE_REGULATION_OF_GENE_EXPRESSION
IMMUNE_EFFECTOR_PROCESS
LYMPHOCYTE_APOPTOTIC_PROCESS
PROTEIN_RNA_COMPLEX_ORGANIZATION
REGULATION_OF_UBIQUITIN_PROTEIN_LIGASE_ACTIVITY
RIBOSOMAL_LARGE_SUBUNIT_BIOGENESIS
REGULATION_OF_UBIQUITIN_PROTEIN_TRANSFERASE_ACTIVI...
RRNA_METABOLIC_PROCESS
GLIAL_CELL_APOPTOTIC_PROCESS
MYELOID_DENDRITIC_CELL_DIFFERENTIATION
REGULATION_OF_SMALL_GTPASE_MEDIATED_SIGNAL_TRANSDU...
REGULATION_OF_MUSCLE_HYPERTROPHY
REGULATION_OF_HOMOTYPIC_CELL_CELL_ADHESION
most shared
most spliced
most unspliced
pathway
NES
0 1 2 3
mode
spliced
unspliced
-log10(padj)
0
2
4
6

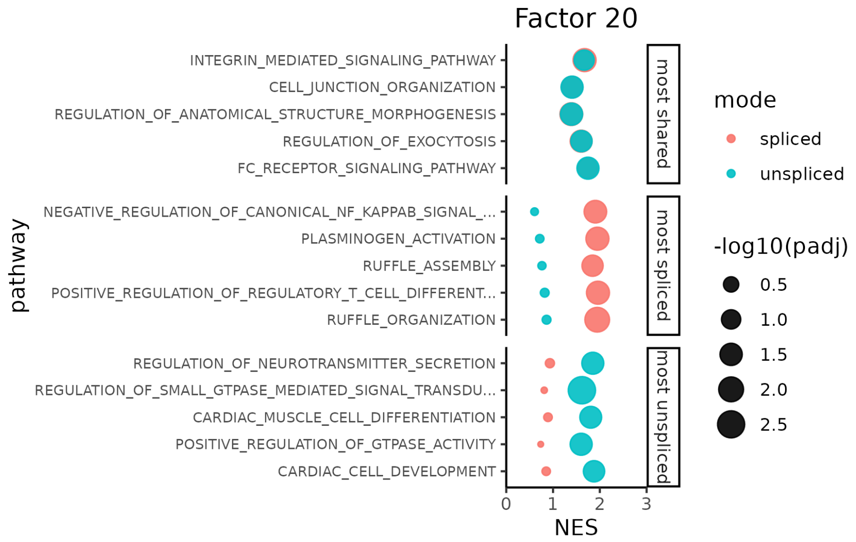
Factor 20
INTEGRIN_MEDIATED_SIGNALING_PATHWAY
CELL_JUNCTION_ORGANIZATION
REGULATION_OF_ANATOMICAL_STRUCTURE_MORPHOGENESIS
REGULATION_OF_EXOCYTOSIS
FC_RECEPTOR_SIGNALING_PATHWAY
NEGATIVE_REGULATION_OF_CANONICAL_NF_KAPPAB_SIGNAL_...
PLASMINOGEN_ACTIVATION
RUFFLE_ASSEMBLY
POSITIVE_REGULATION_OF_REGULATORY_T_CELL_DIFFERENT...
RUFFLE_ORGANIZATION
REGULATION_OF_NEUROTRANSMITTER_SECRETION
REGULATION_OF_SMALL_GTPASE_MEDIATED_SIGNAL_TRANSDU...
CARDIAC_MUSCLE_CELL_DIFFERENTIATION
POSITIVE_REGULATION_OF_GTPASE_ACTIVITY
CARDIAC_CELL_DEVELOPMENT
pathway
most shared
most spliced
most unspliced
0 1 2 3
NES
mode
spliced
unspliced
-log10(padj)
0.5
1.0
1.5
2.0
2.5

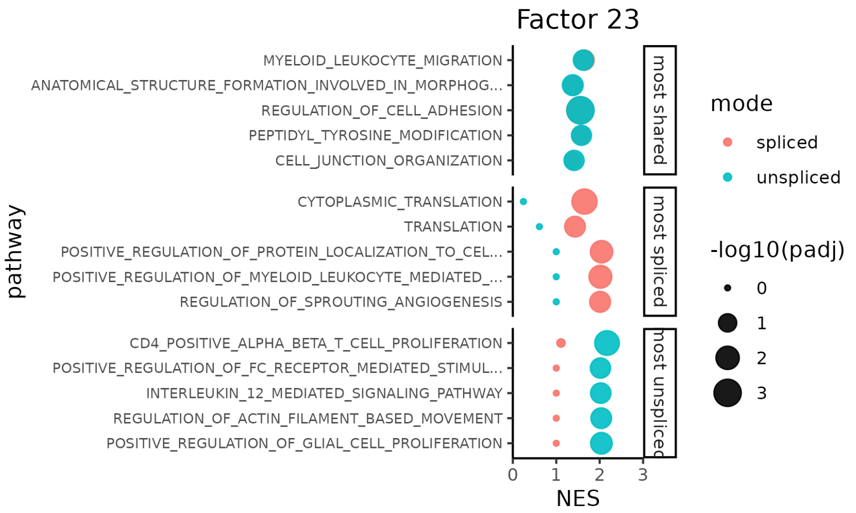
Factor 23
MYELOID_LEUKOCYTE_MIGRATION
ANATOMICAL_STRUCTURE_FORMATION_INVOLVED_IN_MORPHOG...
REGULATION_OF_CELL_ADHESION
PEPTIDYL_TYROSINE_MODIFICATION
CELL_JUNCTION_ORGANIZATION
CYTOPLASMIC_TRANSLATION
TRANSLATION
POSITIVE_REGULATION_OF_PROTEIN_LOCALIZATION_TO_CEL...
POSITIVE_REGULATION_OF_MYELOID_LEUKOCYTE_MEDIATED_...
REGULATION_OF_SPROUTING_ANGIOGENESIS
CD4_POSITIVE_ALPHA_BETA_T_CELL_PROLIFERATION
POSITIVE_REGULATION_OF_FC_RECEPTOR_MEDIATED_STIMUL...
INTERLEUKIN_12_MEDIATED_SIGNALING_PATHWAY
REGULATION_OF_ACTIN_FILAMENT_BASED_MOVEMENT
POSITIVE_REGULATION_OF_GLIAL_CELL_PROLIFERATION
pathway
most shared
most spliced
most unspliced
0 1 2 3
NES
mode
spliced
unspliced
-log10(padj)
0
1
2
3

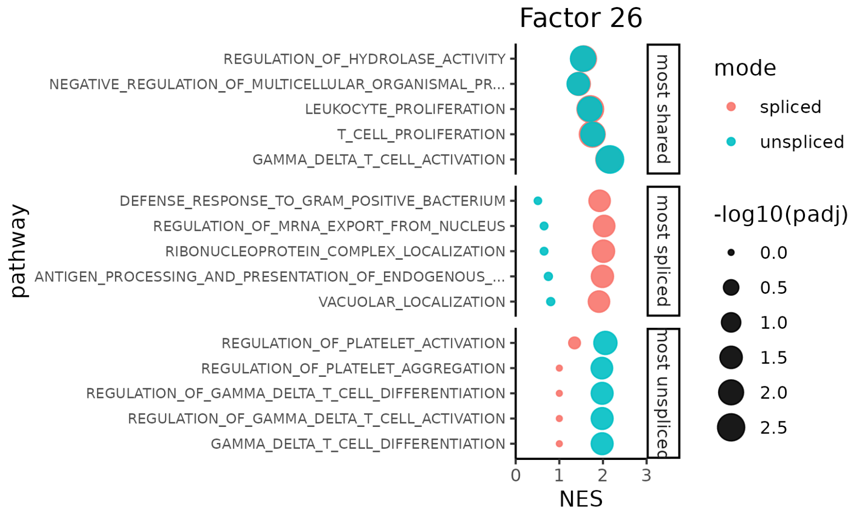
Factor 26
REGULATION_OF_HYDROLASE_ACTIVITY
NEGATIVE_REGULATION_OF_MULTICELLULAR_ORGANISMAL_PR...
LEUKOCYTE_PROLIFERATION
T_CELL_PROLIFERATION
GAMMA_DELTA_T_CELL_ACTIVATION
DEFENSE_RESPONSE_TO_GRAM_POSITIVE_BACTERIUM
REGULATION_OF_MRNA_EXPORT_FROM_NUCLEUS
RIBONUCLEOPROTEIN_COMPLEX_LOCALIZATION
ANTIGEN_PROCESSING_AND_PRESENTATION_OF_ENDOGENOUS_...
VACUOLAR_LOCALIZATION
REGULATION_OF_PLATELET_ACTIVATION
REGULATION_OF_PLATELET_AGGREGATION
REGULATION_OF_GAMMA_DELTA_T_CELL_DIFFERENTIATION
REGULATION_OF_GAMMA_DELTA_T_CELL_ACTIVATION
GAMMA_DELTA_T_CELL_DIFFERENTIATION
pathway
most shared
most spliced
most unspliced
0
1
2
3
NES
mode
spliced
unspliced
-log10(padj)
0.0
0.5
1.0
1.5
2.0
2.5

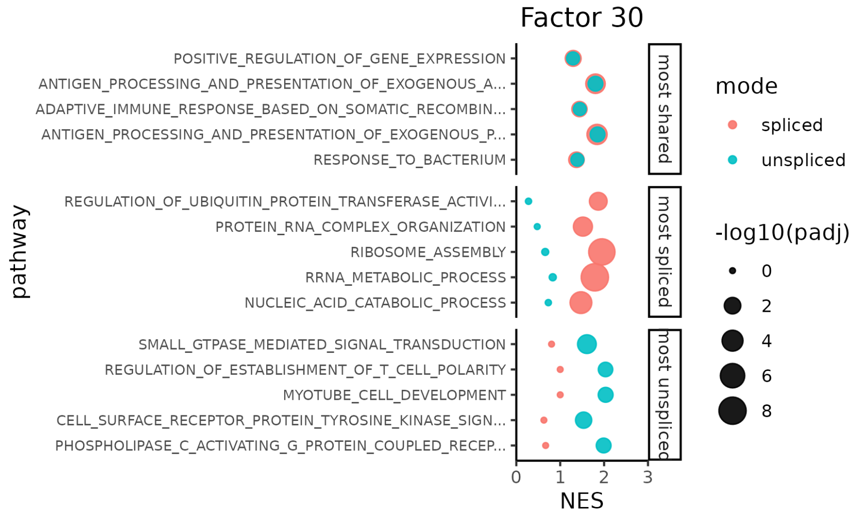
Factor 30
POSITIVE_REGULATION_OF_GENE_EXPRESSION
ANTIGEN_PROCESSING_AND_PRESENTATION_OF_EXOGENOUS_A...
ADAPTIVE_IMMUNE_RESPONSE_BASED_ON_SOMATIC_RECOMBIN...
ANTIGEN_PROCESSING_AND_PRESENTATION_OF_EXOGENOUS_P...
RESPONSE_TO_BACTERIUM
REGULATION_OF_UBIQUITIN_PROTEIN_TRANSFERASE_ACTIVI...
PROTEIN_RNA_COMPLEX_ORGANIZATION
RIBOSOME_ASSEMBLY
RRNA_METABOLIC_PROCESS
NUCLEIC_ACID_CATABOLIC_PROCESS
SMALL_GTPASE_MEDIATED_SIGNAL_TRANSDUCTION
REGULATION_OF_ESTABLISHMENT_OF_T_CELL_POLARITY
MYOTUBE_CELL_DEVELOPMENT
CELL_SURFACE_RECEPTOR_PROTEIN_TYROSINE_KINASE_SIGN...
PHOSPHOLIPASE_C_ACTIVATING_G_PROTEIN_COUPLED_RECEP...
pathway
most shared
most spliced
most unspliced
0
1
2
3
NES
mode
spliced
unspliced
-log10(padj)
0
2
4
6
8

Table 1: **Test Mean Squared Error (MSE) by Modality and Replicate for the Brain dataset.** Each column shows the test reconstruction error from cross-validated NMF across three random replicates per modality. Highlighted values indicate the **best (lowest) test MSE rank** for each replicate, corresponding to the optimal number of latent factors ($k^*$).

| | Added | | | Spliced Only | | | Unspliced Only | | | Concatenated | | |
|---|---|---|---|---|---|---|---|---|---|---|---|---|
| k | Rep 1 | Rep 2 | Rep 3 | Rep 1 | Rep 2 | Rep 3 | Rep 1 | Rep 2 | Rep3 | Rep 1 | Rep 2 | Rep3 |
| 2 | 0.0665 | 0.0664 | 0.0665 | 0.0379 | 0.0379 | 0.0379 | 0.0363 | 0.0362 | 0.0362 | 0.0379 | 0.038 | 0.038 |
| 3 | 0.0646 | 0.0646 | 0.0646 | 0.0373 | 0.0374 | 0.0373 | 0.0352 | 0.0351 | 0.0351 | 0.037 | 0.037 | 0.037 |
| 4 | 0.0631 | 0.0631 | 0.0631 | 0.0369 | 0.0369 | 0.0369 | 0.0345 | 0.0344 | 0.0344 | 0.0363 | 0.0363 | 0.0363 |
| 5 | 0.0618 | 0.0617 | 0.0617 | 0.0366 | 0.0366 | 0.0365 | 0.0341 | 0.034 | 0.0341 | 0.0356 | 0.0356 | 0.0356 |
| 6 | 0.0613 | 0.0612 | 0.0613 | 0.0363 | 0.0363 | 0.0363 | 0.0339 | 0.0338 | 0.0338 | 0.0354 | 0.0354 | 0.0354 |
| 7 | 0.0608 | 0.0607 | 0.0609 | 0.036 | 0.036 | 0.036 | 0.0337 | 0.0335 | 0.0336 | 0.0352 | 0.0352 | 0.0352 |
| 8 | 0.0605 | 0.0603 | 0.0605 | 0.0358 | 0.0358 | 0.0358 | 0.0335 | 0.0334 | 0.0334 | 0.035 | 0.035 | 0.035 |
| 9 | 0.0601 | 0.0601 | 0.0601 | 0.0356 | 0.0357 | 0.0357 | 0.0333 | 0.0332 | 0.0333 | 0.0349 | 0.0348 | 0.0349 |
| 10 | 0.0598 | 0.0597 | 0.0598 | 0.0355 | 0.0355 | 0.0355 | 0.0332 | 0.0331 | 0.0331 | 0.0347 | 0.0347 | 0.0347 |
| 11 | 0.0596 | 0.0595 | 0.0595 | 0.0354 | 0.0354 | 0.0354 | 0.0331 | 0.033 | 0.033 | 0.0346 | 0.0345 | 0.0346 |
| 12 | 0.0594 | 0.0593 | 0.0593 | 0.0354 | 0.0353 | 0.0354 | 0.0329 | 0.0329 | 0.0329 | 0.0344 | 0.0344 | 0.0344 |
| 13 | 0.0592 | 0.059 | 0.0591 | 0.0353 | 0.0353 | 0.0353 | 0.0328 | 0.0328 | 0.0328 | 0.0343 | 0.0343 | 0.0343 |
| 14 | 0.059 | 0.0589 | 0.059 | 0.0352 | 0.0352 | 0.0352 | 0.0328 | 0.0327 | 0.0327 | 0.0342 | 0.0342 | 0.0342 |
| 15 | 0.0588 | 0.0587 | 0.0588 | 0.0352 | 0.0352 | 0.0351 | 0.0327 | 0.0326 | 0.0326 | 0.0341 | 0.0341 | 0.0342 |
| 16 | 0.0587 | 0.0586 | 0.0586 | 0.0351 | 0.0351 | 0.0351 | 0.0326 | 0.0325 | 0.0325 | 0.0341 | 0.034 | 0.0341 |
| 17 | 0.0586 | 0.0585 | 0.0586 | 0.035 | 0.035 | 0.035 | 0.0325 | 0.0324 | 0.0324 | 0.034 | 0.034 | 0.034 |
| 18 | 0.0585 | 0.0584 | 0.0584 | 0.035 | 0.035 | 0.035 | 0.0324 | 0.0324 | 0.0324 | 0.0339 | 0.0339 | 0.034 |
| 19 | 0.0583 | 0.0583 | 0.0583 | 0.0349 | 0.0349 | 0.0349 | 0.0324 | 0.0323 | 0.0323 | 0.0339 | 0.0339 | 0.0339 |
| 20 | 0.0583 | 0.0582 | 0.0582 | 0.0349 | 0.0349 | 0.0349 | 0.0323 | 0.0323 | 0.0323 | 0.0338 | 0.0338 | 0.0339 |
| 21 | 0.0582 | 0.058 | 0.0581 | 0.0348 | 0.0348 | 0.0349 | 0.0323 | 0.0322 | 0.0322 | 0.0338 | 0.0338 | 0.0338 |
| 22 | 0.058 | 0.0579 | 0.058 | 0.0348 | 0.0348 | 0.0356 | 0.0322 | 0.0321 | 0.0322 | 0.0337 | 0.0337 | 0.0337 |
| 23 | 0.058 | 0.0578 | 0.058 | 0.0348 | 0.0348 | 0.0348 | 0.0322 | 0.0321 | 0.0321 | 0.0337 | 0.0337 | 0.0337 |
| 24 | 0.0579 | 0.0578 | 0.0579 | 0.0348 | 0.0348 | 0.0348 | 0.0322 | 0.0321 | 0.0321 | 0.0336 | 0.0336 | 0.0337 |
| 25 | 0.0578 | 0.0577 | 0.0578 | 0.0349 | 0.0347 | 0.0348 | 0.0321 | 0.032 | 0.032 | 0.0336 | 0.0336 | 0.0336 |
| 26 | 0.0577 | 0.0576 | 0.0577 | 0.0349 | 0.0347 | 0.0348 | 0.0321 | 0.032 | 0.032 | 0.0336 | 0.0336 | 0.0336 |
| 27 | 0.0577 | 0.0576 | 0.0576 | 0.0483 | 0.0347 | 0.0347 | 0.0321 | 0.032 | 0.032 | 0.0335 | 0.0335 | 0.0336 |
| 28 | 0.0576 | 0.0575 | 0.0575 | 0.0357 | 0.0348 | 0.0347 | 0.032 | 0.032 | 0.0319 | 0.0335 | 0.0335 | 0.0335 |
| 29 | 0.0575 | 0.0575 | 0.0575 | 0.0503 | 0.0347 | 0.0347 | 0.032 | 0.0319 | 0.0319 | 0.0335 | 0.0334 | 0.0335 |
| 30 | 0.0575 | 0.0574 | 0.0575 | 0.0435 | 0.0347 | 0.0347 | 0.032 | 0.0319 | 0.0319 | 0.0334 | 0.0334 | 0.0335 |
| 31 | 0.0574 | 0.0573 | 0.0574 | 0.0415 | 0.0397 | 0.035 | 0.032 | 0.0319 | 0.0319 | 0.0334 | 0.0334 | 0.0334 |
| 32 | 0.0574 | 0.0573 | 0.0573 | 0.0575 | 0.0356 | 0.035 | 0.0319 | 0.0319 | 0.0319 | 0.0334 | 0.0334 | 0.0334 |
| 33 | 0.0573 | 0.0573 | 0.0573 | 0.0659 | 0.035 | 0.0347 | 0.0319 | 0.0319 | 0.0319 | 0.0334 | 0.0333 | 0.0334 |
| 34 | 0.0573 | 0.0572 | 0.0573 | 0.0578 | 0.0352 | 0.0347 | 0.0319 | 0.0318 | 0.0319 | 0.0333 | 0.0333 | 0.0334 |
| 35 | 0.0572 | 0.0572 | 0.0572 | 0.0652 | 0.0361 | 0.0352 | 0.0319 | 0.0318 | 0.0318 | 0.0333 | 0.0333 | 0.0333 |
| 36 | 0.0572 | 0.0571 | 0.0572 | 0.0962 | 0.0391 | 0.037 | 0.0319 | 0.0318 | 0.0318 | 0.0333 | 0.0333 | 0.0333 |
| 37 | 0.0572 | 0.0571 | 0.0572 | 0.1084 | 0.0403 | 0.0379 | 0.0319 | 0.0318 | 0.0318 | 0.0333 | 0.0333 | 0.0333 |
| 38 | 0.0571 | 0.0571 | 0.0571 | 0.1036 | 0.0591 | 0.0411 | 0.0369 | 0.0318 | 0.0318 | 0.0332 | 0.0332 | 0.0333 |
| 39 | 0.0571 | 0.0571 | 0.0571 | 0.1243 | 0.0837 | 0.0482 | 0.0345 | 0.0318 | 0.0318 | 0.0332 | 0.0332 | 0.0333 |
| 40 | 0.0571 | 0.057 | 0.057 | 0.1052 | 0.088 | 0.0685 | 0.0352 | 0.0318 | 0.0318 | 0.0332 | 0.0332 | 0.0332 |
| 41 | 0.0583 | 0.057 | 0.057 | 0.116 | 0.0876 | 0.0739 | 0.0466 | 0.0318 | 0.0319 | 0.0332 | 0.0332 | 0.0332 |
| 42 | 0.0587 | 0.0572 | 0.057 | 0.1009 | 0.0876 | 0.0712 | 0.0394 | 0.0318 | 0.0561 | 0.0345 | 0.0332 | 0.0332 |
| 43 | 0.058 | 0.057 | 0.057 | 0.1334 | 0.0965 | 0.0974 | 0.0419 | 0.0551 | 0.0622 | 0.0434 | 0.0332 | 0.0332 |
| 44 | 0.0701 | 0.0577 | 0.0571 | 0.1391 | 0.078 | 0.0956 | 0.0326 | 0.0592 | 0.0421 | 0.0405 | 0.0332 | 0.0332 |
| 45 | 0.0572 | 0.0577 | 0.057 | 0.1783 | 0.0846 | 0.0715 | 0.0345 | 0.0789 | 0.0522 | 0.0417 | 0.0333 | 0.0332 |
| 46 | 0.0585 | 0.0609 | 0.0574 | 0.1641 | 0.0836 | 0.1115 | 0.043 | 0.0733 | 0.0482 | 0.0407 | 0.0335 | 0.0388 |
| 47 | 0.0667 | 0.0647 | 0.0667 | 0.1545 | 0.0971 | 0.1363 | 0.0518 | 0.0733 | 0.0404 | 0.0415 | 0.0344 | 0.0332 |
| 48 | 0.0696 | 0.0643 | 0.0654 | 0.1643 | 0.1186 | 0.139 | 0.0518 | 0.077 | 0.0439 | 0.0455 | 0.0357 | 0.0411 |
| 49 | 0.066 | 0.0786 | 0.067 | 0.1887 | 0.2129 | 0.0899 | 0.0633 | 0.075 | 0.05 | 0.0458 | 0.0332 | 0.0393 |
| 50 | 0.0747 | 0.0588 | 0.076 | 0.1663 | 0.2396 | 0.1056 | 0.0798 | 0.0794 | 0.0535 | 0.0332 | 0.0331 | 0.0356 |
| 51 | 0.0743 | 0.0713 | 0.074 | 0.1927 | 0.2528 | 0.1255 | 0.0704 | 0.0905 | 0.0612 | 0.0349 | 0.0332 | 0.0467 |
| 52 | 0.0747 | 0.0711 | 0.0877 | 0.2261 | 0.235 | 0.1479 | 0.0736 | 0.1111 | 0.0589 | 0.0498 | 0.0338 | 0.0539 |
| 53 | 0.0674 | 0.0762 | 0.0696 | 0.2415 | 0.2687 | 0.1884 | 0.0942 | 0.1595 | 0.0796 | 0.0587 | 0.0385 | 0.0565 |
| 54 | 0.0737 | 0.103 | 0.0849 | 0.3011 | 0.2983 | 0.1871 | 0.1307 | 0.1982 | 0.0627 | 0.0844 | 0.047 | 0.0518 |
| 55 | 0.0787 | 0.1104 | 0.0778 | 0.2892 | 0.3059 | 0.1814 | 0.1131 | 0.2378 | 0.1151 | 0.1071 | 0.0349 | 0.0626 |
| 56 | 0.0827 | 0.0998 | 0.1003 | 0.2963 | 0.3121 | 0.2033 | 0.1078 | 0.2527 | 0.1194 | 0.1195 | 0.0337 | 0.0613 |
| 57 | 0.0796 | 0.1066 | 0.0934 | 0.3356 | 0.3203 | 0.2273 | 0.1761 | 0.2756 | 0.1474 | 0.1053 | 0.0345 | 0.0917 |
| 58 | 0.075 | 0.1296 | 0.0941 | 0.3602 | 0.351 | 0.2872 | 0.1543 | 0.2324 | 0.1311 | 0.1019 | 0.0374 | 0.0917 |
| 59 | 0.0737 | 0.1337 | 0.0912 | 0.3601 | 0.3413 | 0.3492 | 0.1637 | 0.2469 | 0.1416 | 0.1014 | 0.0368 | 0.0968 |
| 60 | 0.1002 | 0.1199 | 0.0976 | 0.407 | 0.3776 | 0.3546 | 0.2 | 0.3066 | 0.1736 | 0.1362 | 0.0367 | 0.0938 |
| 61 | 0.0991 | 0.138 | 0.1088 | 0.3893 | 0.4061 | 0.4038 | 0.2465 | 0.4119 | 0.2006 | 0.1432 | 0.0359 | 0.1039 |
| 62 | 0.1148 | 0.1491 | 0.1228 | 0.4422 | 0.4549 | 0.4814 | 0.2222 | 0.4604 | 0.2618 | 0.1398 | 0.0406 | 0.1038 |
| 63 | 0.0954 | 0.1507 | 0.131 | 0.4691 | 0.4724 | 0.4814 | 0.2626 | 0.5093 | 0.2952 | 0.1478 | 0.0412 | 0.1267 |
| 64 | 0.1225 | 0.1604 | 0.1327 | 0.5866 | 0.5232 | 0.4902 | 0.2645 | 0.572 | 0.3027 | 0.1691 | 0.0494 | 0.1309 |
| 65 | 0.1315 | 0.1541 | 0.1365 | 0.6227 | 0.5607 | 0.5651 | 0.3422 | 0.5363 | 0.3121 | 0.167 | 0.0612 | 0.1316 |
| 66 | 0.1267 | 0.162 | 0.1458 | 0.6833 | 0.6158 | 0.6281 | 0.3223 | 0.6858 | 0.3657 | 0.1764 | 0.069 | 0.1022 |
| 67 | 0.1345 | 0.1851 | 0.1386 | 0.7035 | 0.6472 | 0.6208 | 0.352 | 0.6866 | 0.3542 | 0.1679 | 0.0646 | 0.1605 |
| 68 | 0.1403 | 0.1908 | 0.1489 | 0.7165 | 0.7346 | 0.6902 | 0.4498 | 0.7583 | 0.3712 | 0.178 | 0.0899 | 0.1685 |
| 69 | 0.1434 | 0.2133 | 0.1245 | 0.8113 | 0.8543 | 0.735 | 0.4824 | 0.7566 | 0.385 | 0.1894 | 0.1157 | 0.1768 |
| 70 | 0.1591 | 0.21 | 0.1803 | 0.9153 | 0.8849 | 0.7732 | 0.5266 | 0.8035 | 0.4939 | 0.1938 | 0.0884 | 0.1858 |
| 71 | 0.1351 | 0.2214 | 0.1808 | 1.0455 | 0.8834 | 0.8488 | 0.5426 | 0.8355 | 0.5174 | 0.1945 | 0.1096 | 0.1959 |
| 72 | 0.197 | 0.2071 | 0.1859 | 1.0713 | 0.9508 | 0.8447 | 0.567 | 0.9225 | 0.6162 | 0.1986 | 0.1053 | 0.1906 |
| 73 | 0.2012 | 0.2062 | 0.2502 | 1.1472 | 0.9871 | 0.9337 | 0.5691 | 0.9747 | 0.5791 | 0.1819 | 0.1068 | 0.1879 |
| 74 | 0.1994 | 0.2501 | 0.2782 | 1.2015 | 1.1517 | 1.0124 | 0.6385 | 1.0345 | 0.662 | 0.207 | 0.1437 | 0.2312 |
| 75 | 0.1928 | 0.2403 | 0.3383 | 1.2573 | 1.1944 | 1.081 | 0.7996 | 1.0208 | 0.7081 | 0.1844 | 0.149 | 0.2281 |
| 76 | 0.3012 | 0.2907 | 0.3975 | 1.3221 | 1.2336 | 1.1358 | 0.8148 | 1.3043 | 0.8423 | 0.195 | 0.1364 | 0.249 |
| 77 | 0.3254 | 0.3836 | 0.3888 | 1.3812 | 1.2569 | 1.1384 | 0.8355 | 1.2526 | 0.8221 | 0.1924 | 0.1267 | 0.2632 |
| 78 | 0.3625 | 0.3943 | 0.2948 | 1.4708 | 1.3041 | 1.2114 | 0.8628 | 1.2614 | 0.905 | 0.2174 | 0.1511 | 0.2834 |
| 79 | 0.4081 | 0.4024 | 0.2123 | 1.5062 | 1.4751 | 1.2719 | 0.9255 | 1.4084 | 1.0022 | 0.2659 | 0.1834 | 0.2859 |
| 80 | 0.4268 | 0.4186 | 0.2458 | 1.542 | 1.5874 | 1.3208 | 0.9203 | 1.5088 | 1.0206 | 0.2681 | 0.2171 | 0.3134 |

Table 2: **Test Mean Squared Error (MSE) by Modality and Replicate for the Heart dataset.** Each column shows the test reconstruction error from cross-validated NMF across three random replicates per modality. Highlighted values indicate the **best (lowest) test MSE rank** for each replicate, corresponding to the optimal number of latent factors ($k^*$).

| | Added | | | Spliced Only | | | Unspliced Only | | | Concatenated | | |
|---|---|---|---|---|---|---|---|---|---|---|---|---|
| k | Rep 1 | Rep 2 | Rep 3 | Rep 1 | Rep 2 | Rep 3 | Rep 1 | Rep 2 | Rep3 | Rep 1 | Rep 2 | Rep3 |
| 2 | 0.076 | 0.0762 | 0.0762 | 0.0393 | 0.0392 | 0.0393 | 0.0433 | 0.0433 | 0.0434 | 0.0426 | 0.0426 | 0.0426 |
| 3 | 0.0726 | 0.0727 | 0.0727 | 0.0376 | 0.0377 | 0.0376 | 0.0426 | 0.0426 | 0.0426 | 0.0408 | 0.0408 | 0.0408 |
| 4 | 0.0711 | 0.0712 | 0.0712 | 0.0365 | 0.0364 | 0.0364 | 0.042 | 0.042 | 0.0421 | 0.0401 | 0.0401 | 0.0401 |
| 5 | 0.0697 | 0.0701 | 0.0701 | 0.0358 | 0.0358 | 0.0358 | 0.0417 | 0.0416 | 0.0417 | 0.0392 | 0.0392 | 0.0392 |
| 6 | 0.0687 | 0.0688 | 0.0688 | 0.0353 | 0.0353 | 0.0353 | 0.0414 | 0.0414 | 0.0414 | 0.0389 | 0.0389 | 0.0389 |
| 7 | 0.0681 | 0.0681 | 0.0681 | 0.0348 | 0.0349 | 0.0348 | 0.0411 | 0.0411 | 0.0412 | 0.0385 | 0.0385 | 0.0385 |
| 8 | 0.0675 | 0.0674 | 0.0674 | 0.0346 | 0.0346 | 0.0346 | 0.0409 | 0.0409 | 0.041 | 0.0383 | 0.0382 | 0.0383 |
| 9 | 0.067 | 0.0671 | 0.0671 | 0.0343 | 0.0343 | 0.0343 | 0.0407 | 0.0407 | 0.0408 | 0.038 | 0.0381 | 0.038 |
| 10 | 0.0666 | 0.0667 | 0.0667 | 0.034 | 0.0341 | 0.0341 | 0.0405 | 0.0405 | 0.0406 | 0.0378 | 0.0378 | 0.0379 |
| 11 | 0.0662 | 0.0663 | 0.0663 | 0.0339 | 0.0339 | 0.0339 | 0.0404 | 0.0404 | 0.0405 | 0.0376 | 0.0376 | 0.0376 |
| 12 | 0.0659 | 0.066 | 0.066 | 0.0337 | 0.0337 | 0.0337 | 0.0403 | 0.0403 | 0.0404 | 0.0375 | 0.0375 | 0.0375 |
| 13 | 0.0656 | 0.0656 | 0.0656 | 0.0335 | 0.0335 | 0.0336 | 0.0402 | 0.0402 | 0.0402 | 0.0373 | 0.0373 | 0.0373 |
| 14 | 0.0653 | 0.0654 | 0.0654 | 0.0334 | 0.0334 | 0.0334 | 0.0401 | 0.0401 | 0.0402 | 0.0372 | 0.0372 | 0.0372 |
| 15 | 0.0651 | 0.0651 | 0.0651 | 0.0332 | 0.0332 | 0.0332 | 0.0401 | 0.04 | 0.0401 | 0.037 | 0.037 | 0.0371 |
| 16 | 0.0649 | 0.0648 | 0.0648 | 0.0331 | 0.0331 | 0.0331 | 0.04 | 0.04 | 0.04 | 0.037 | 0.037 | 0.0369 |
| 17 | 0.0647 | 0.0646 | 0.0646 | 0.033 | 0.033 | 0.033 | 0.0399 | 0.0399 | 0.04 | 0.0369 | 0.0368 | 0.0368 |
| 18 | 0.0645 | 0.0646 | 0.0646 | 0.0329 | 0.0329 | 0.0329 | 0.0399 | 0.0399 | 0.0399 | 0.0368 | 0.0368 | 0.0367 |
| 19 | 0.0644 | 0.0644 | 0.0644 | 0.0328 | 0.0328 | 0.0328 | 0.0399 | 0.0398 | 0.0399 | 0.0367 | 0.0367 | 0.0367 |
| 20 | 0.0642 | 0.0643 | 0.0643 | 0.0328 | 0.0328 | 0.0328 | 0.0398 | 0.0398 | 0.0399 | 0.0366 | 0.0366 | 0.0366 |
| 21 | 0.0642 | 0.0642 | 0.0642 | 0.0327 | 0.0327 | 0.0327 | 0.0398 | 0.0398 | 0.0398 | 0.0366 | 0.0365 | 0.0365 |
| 22 | 0.0641 | 0.0641 | 0.0641 | 0.0327 | 0.0327 | 0.0326 | 0.0398 | 0.0398 | 0.0398 | 0.0365 | 0.0365 | 0.0365 |
| 23 | 0.0639 | 0.064 | 0.064 | 0.0326 | 0.0326 | 0.0326 | 0.0397 | 0.0397 | 0.0398 | 0.0365 | 0.0364 | 0.0364 |
| 24 | 0.0638 | 0.0639 | 0.0639 | 0.0327 | 0.0325 | 0.0325 | 0.0397 | 0.0397 | 0.0398 | 0.0364 | 0.0364 | 0.0364 |
| 25 | 0.0638 | 0.0638 | 0.0638 | 0.0325 | 0.0325 | 0.0325 | 0.0397 | 0.0397 | 0.0398 | 0.0363 | 0.0363 | 0.0363 |
| 26 | 0.0637 | 0.0637 | 0.0637 | 0.0324 | 0.0325 | 0.0324 | 0.0397 | 0.0397 | 0.0398 | 0.0363 | 0.0363 | 0.0363 |
| 27 | 0.0636 | 0.0636 | 0.0636 | 0.0324 | 0.0324 | 0.0324 | 0.0397 | 0.0397 | 0.0397 | 0.0363 | 0.0363 | 0.0363 |
| 28 | 0.0635 | 0.0636 | 0.0636 | 0.0324 | 0.0324 | 0.0324 | 0.0397 | 0.0397 | 0.0397 | 0.0362 | 0.0362 | 0.0362 |
| 29 | 0.0635 | 0.0635 | 0.0635 | 0.0324 | 0.0323 | 0.0323 | 0.0396 | 0.0397 | 0.0397 | 0.0362 | 0.0362 | 0.0362 |
| 30 | 0.0634 | 0.0635 | 0.0635 | 0.0323 | 0.0324 | 0.0324 | 0.0396 | 0.0396 | 0.0397 | 0.0362 | 0.0362 | 0.0362 |
| 31 | 0.0634 | 0.0634 | 0.0634 | 0.0323 | 0.0323 | 0.0329 | 0.0396 | 0.0396 | 0.0397 | 0.0361 | 0.0361 | 0.0361 |
| 32 | 0.0633 | 0.0634 | 0.0634 | 0.0323 | 0.0325 | 0.0325 | 0.0396 | 0.0396 | 0.0397 | 0.0361 | 0.0361 | 0.0361 |
| 33 | 0.0633 | 0.0633 | 0.0633 | 0.0323 | 0.0323 | 0.0334 | 0.0396 | 0.0396 | 0.0397 | 0.0361 | 0.0361 | 0.0361 |
| 34 | 0.0632 | 0.0632 | 0.0632 | 0.0323 | 0.0325 | 0.0324 | 0.0396 | 0.0396 | 0.0397 | 0.036 | 0.0361 | 0.0361 |
| 35 | 0.0632 | 0.0632 | 0.0632 | 0.0386 | 0.035 | 0.0437 | 0.0396 | 0.0396 | 0.0397 | 0.036 | 0.036 | 0.036 |
| 36 | 0.0632 | 0.0631 | 0.0631 | 0.0416 | 0.0404 | 0.0474 | 0.0396 | 0.0396 | 0.0397 | 0.036 | 0.036 | 0.036 |
| 37 | 0.0631 | 0.0631 | 0.0631 | 0.0461 | 0.0322 | 0.0554 | 0.0396 | 0.0396 | 0.0397 | 0.036 | 0.036 | 0.036 |
| 38 | 0.0631 | 0.0631 | 0.0631 | 0.0889 | 0.034 | 0.062 | 0.0396 | 0.0396 | 0.0397 | 0.036 | 0.036 | 0.036 |
| 39 | 0.063 | 0.063 | 0.063 | 0.0655 | 0.0341 | 0.0559 | 0.0396 | 0.0396 | 0.0397 | 0.036 | 0.0359 | 0.036 |
| 40 | 0.063 | 0.063 | 0.063 | 0.051 | 0.0364 | 0.0449 | 0.0403 | 0.0396 | 0.0397 | 0.0359 | 0.0359 | 0.0359 |
| 41 | 0.063 | 0.063 | 0.063 | 0.0531 | 0.0446 | 0.0468 | 0.0436 | 0.0396 | 0.0479 | 0.0359 | 0.0359 | 0.0376 |
| 42 | 0.0629 | 0.063 | 0.063 | 0.0903 | 0.05 | 0.0492 | 0.0505 | 0.0397 | 0.0458 | 0.0359 | 0.0386 | 0.0359 |
| 43 | 0.0629 | 0.0629 | 0.0629 | 0.0981 | 0.0522 | 0.0598 | 0.0556 | 0.0397 | 0.0397 | 0.0359 | 0.0359 | 0.0365 |
| 44 | 0.063 | 0.0629 | 0.0629 | 0.0558 | 0.0468 | 0.0585 | 0.0571 | 0.0397 | 0.0432 | 0.0359 | 0.0359 | 0.0359 |
| 45 | 0.0682 | 0.0629 | 0.0629 | 0.0533 | 0.0695 | 0.0606 | 0.0509 | 0.0397 | 0.0397 | 0.0359 | 0.0359 | 0.0359 |
| 46 | 0.0629 | 0.0629 | 0.0629 | 0.0629 | 0.0624 | 0.0612 | 0.0449 | 0.0397 | 0.0397 | 0.0359 | 0.0359 | 0.0359 |
| 47 | 0.0629 | 0.0629 | 0.0629 | 0.0853 | 0.0735 | 0.0769 | 0.0502 | 0.0397 | 0.0397 | 0.0358 | 0.0358 | 0.0358 |
| 48 | 0.0629 | 0.0629 | 0.0629 | 0.0861 | 0.0654 | 0.0712 | 0.056 | 0.0397 | 0.0397 | 0.0358 | 0.0359 | 0.0358 |
| 49 | 0.0629 | 0.0629 | 0.0629 | 0.091 | 0.1649 | 0.0788 | 0.0574 | 0.0397 | 0.0479 | 0.0358 | 0.04 | 0.0358 |
| 50 | 0.0629 | 0.0629 | 0.0629 | 0.0987 | 0.1703 | 0.0727 | 0.0568 | 0.0397 | 0.049 | 0.0358 | 0.0376 | 0.0358 |
| 51 | 0.0629 | 0.0629 | 0.0629 | 0.0913 | 0.1668 | 0.0891 | 0.0589 | 0.0397 | 0.0439 | 0.0391 | 0.0358 | 0.0358 |
| 52 | 0.0629 | 0.0629 | 0.0629 | 0.1032 | 0.1103 | 0.0905 | 0.0583 | 0.0401 | 0.0397 | 0.0378 | 0.0358 | 0.0358 |
| 53 | 0.0629 | 0.0628 | 0.0628 | 0.1105 | 0.16 | 0.0902 | 0.0592 | 0.0397 | 0.0398 | 0.0395 | 0.0358 | 0.0436 |
| 54 | 0.0633 | 0.0649 | 0.0649 | 0.1099 | 0.1642 | 0.1075 | 0.0603 | 0.0397 | 0.0398 | 0.0398 | 0.0358 | 0.0396 |
| 55 | 0.088 | 0.0638 | 0.0638 | 0.1336 | 0.2223 | 0.1583 | 0.0599 | 0.0398 | 0.0453 | 0.0375 | 0.0374 | 0.0453 |
| 56 | 0.0815 | 0.0645 | 0.0645 | 0.1299 | 0.2229 | 0.1572 | 0.0629 | 0.0456 | 0.0593 | 0.0359 | 0.0376 | 0.0412 |
| 57 | 0.0885 | 0.0669 | 0.0669 | 0.1081 | 0.2198 | 0.1612 | 0.0605 | 0.0464 | 0.0464 | 0.036 | 0.0358 | 0.0406 |
| 58 | 0.1029 | 0.0696 | 0.0696 | 0.0937 | 0.2367 | 0.1697 | 0.0513 | 0.0471 | 0.05 | 0.0367 | 0.0374 | 0.0431 |
| 59 | 0.1038 | 0.0687 | 0.0687 | 0.1279 | 0.177 | 0.1711 | 0.0499 | 0.0463 | 0.0449 | 0.0367 | 0.0438 | 0.0365 |
| 60 | 0.1129 | 0.063 | 0.063 | 0.106 | 0.2211 | 0.1594 | 0.0546 | 0.0496 | 0.0466 | 0.036 | 0.0422 | 0.0374 |
| 61 | 0.1113 | 0.066 | 0.066 | 0.1693 | 0.196 | 0.1853 | 0.0605 | 0.0526 | 0.0579 | 0.0379 | 0.0465 | 0.0476 |
| 62 | 0.1082 | 0.0671 | 0.0671 | 0.1759 | 0.2124 | 0.1884 | 0.0547 | 0.0559 | 0.0607 | 0.0394 | 0.0464 | 0.0453 |
| 63 | 0.1049 | 0.07 | 0.07 | 0.1758 | 0.17 | 0.176 | 0.0587 | 0.0649 | 0.0469 | 0.063 | 0.0465 | 0.0428 |
| 64 | 0.1056 | 0.0745 | 0.0745 | 0.1966 | 0.1962 | 0.1776 | 0.0748 | 0.0646 | 0.0559 | 0.0658 | 0.0487 | 0.0425 |
| 65 | 0.1092 | 0.0731 | 0.0731 | 0.2018 | 0.2046 | 0.1722 | 0.0803 | 0.0696 | 0.0527 | 0.0683 | 0.049 | 0.0411 |
| 66 | 0.0881 | 0.063 | 0.063 | 0.247 | 0.2372 | 0.2123 | 0.164 | 0.0808 | 0.058 | 0.0711 | 0.05 | 0.0449 |
| 67 | 0.0911 | 0.1119 | 0.1119 | 0.2339 | 0.2504 | 0.1857 | 0.1507 | 0.084 | 0.0731 | 0.0747 | 0.051 | 0.0499 |
| 68 | 0.0854 | 0.1037 | 0.1037 | 0.2115 | 0.2391 | 0.2132 | 0.1758 | 0.0863 | 0.0932 | 0.0679 | 0.077 | 0.0498 |
| 69 | 0.0863 | 0.1052 | 0.1052 | 0.2493 | 0.259 | 0.2327 | 0.189 | 0.0979 | 0.0764 | 0.0708 | 0.0769 | 0.0503 |
| 70 | 0.0845 | 0.1085 | 0.1085 | 0.2407 | 0.2892 | 0.3097 | 0.1969 | 0.1202 | 0.0808 | 0.0693 | 0.083 | 0.0563 |
| 71 | 0.0901 | 0.1074 | 0.1074 | 0.2548 | 0.2969 | 0.3215 | 0.2128 | 0.1547 | 0.1057 | 0.0716 | 0.0773 | 0.0582 |
| 72 | 0.0909 | 0.1092 | 0.1092 | 0.2361 | 0.3212 | 0.3309 | 0.2948 | 0.1693 | 0.1146 | 0.0677 | 0.087 | 0.0668 |
| 73 | 0.0927 | 0.1257 | 0.1257 | 0.2483 | 0.3139 | 0.3901 | 0.3309 | 0.1644 | 0.1885 | 0.0687 | 0.0948 | 0.0692 |
| 74 | 0.0876 | 0.1221 | 0.1221 | 0.2668 | 0.3376 | 0.4802 | 0.4081 | 0.1703 | 0.2324 | 0.0706 | 0.0971 | 0.0603 |
| 75 | 0.1069 | 0.1272 | 0.1272 | 0.2901 | 0.4031 | 0.4566 | 0.4669 | 0.2639 | 0.2507 | 0.0837 | 0.0782 | 0.0645 |
| 76 | 0.1079 | 0.125 | 0.125 | 0.2915 | 0.4315 | 0.4979 | 0.4768 | 0.3065 | 0.3471 | 0.0766 | 0.0822 | 0.0627 |
| 77 | 0.1121 | 0.1317 | 0.1317 | 0.3041 | 0.4685 | 0.5273 | 0.5274 | 0.4272 | 0.2739 | 0.0715 | 0.0987 | 0.0642 |
| 78 | 0.1243 | 0.1379 | 0.1379 | 0.3485 | 0.5029 | 0.5909 | 0.5918 | 0.4309 | 0.3612 | 0.0717 | 0.0731 | 0.0658 |
| 79 | 0.1201 | 0.139 | 0.139 | 0.4488 | 0.6037 | 0.5978 | 0.6772 | 0.4351 | 0.4252 | 0.0706 | 0.0771 | 0.0724 |
| 80 | 0.1254 | 0.1341 | 0.1341 | 0.4401 | 0.6237 | 0.6465 | 0.6937 | 0.4633 | 0.4423 | 0.0776 | 0.0733 | 0.0705 |

Table 3: **Test Mean Squared Error (MSE) by Modality and Replicate for the Neuron dataset.** Each column shows the test reconstruction error from cross-validated NMF across three random replicates per modality. Highlighted values indicate the **best (lowest) test MSE rank** for each replicate, corresponding to the optimal number of latent factors ($k^*$).

| | Added | | | Spliced Only | | | Unspliced Only | | | Concatenated | | |
|---|---|---|---|---|---|---|---|---|---|---|---|---|
| k | Rep 1 | Rep 2 | Rep 3 | Rep 1 | Rep 2 | Rep 3 | Rep 1 | Rep 2 | Rep3 | Rep 1 | Rep 2 | Rep3 |
| 2 | 0.0717 | 0.0717 | 0.0717 | 0.0427 | 0.0428 | 0.0428 | 0.0368 | 0.0368 | 0.0368 | 0.0405 | 0.0406 | 0.0406 |
| 3 | 0.0698 | 0.0699 | 0.0698 | 0.042 | 0.0421 | 0.0421 | 0.0357 | 0.0357 | 0.0358 | 0.0396 | 0.0397 | 0.0397 |
| 4 | 0.0684 | 0.0684 | 0.0683 | 0.0415 | 0.0416 | 0.0416 | 0.0351 | 0.0351 | 0.0351 | 0.0389 | 0.0389 | 0.0389 |
| 5 | 0.0671 | 0.0671 | 0.0671 | 0.0411 | 0.0412 | 0.0412 | 0.0347 | 0.0347 | 0.0347 | 0.0383 | 0.0383 | 0.0383 |
| 6 | 0.0665 | 0.0665 | 0.0664 | 0.0408 | 0.0409 | 0.0409 | 0.0345 | 0.0345 | 0.0345 | 0.038 | 0.038 | 0.0381 |
| 7 | 0.0661 | 0.0662 | 0.0661 | 0.0406 | 0.0407 | 0.0407 | 0.0342 | 0.0342 | 0.0343 | 0.0378 | 0.0378 | 0.0378 |
| 8 | 0.0657 | 0.0657 | 0.0658 | 0.0404 | 0.0405 | 0.0405 | 0.0341 | 0.0341 | 0.0341 | 0.0377 | 0.0377 | 0.0377 |
| 9 | 0.0654 | 0.0654 | 0.0654 | 0.0402 | 0.0403 | 0.0403 | 0.0339 | 0.0339 | 0.034 | 0.0375 | 0.0375 | 0.0375 |
| 10 | 0.065 | 0.0651 | 0.0651 | 0.0401 | 0.0402 | 0.0401 | 0.0338 | 0.0338 | 0.0338 | 0.0373 | 0.0373 | 0.0374 |
| 11 | 0.0648 | 0.0648 | 0.0648 | 0.04 | 0.0401 | 0.04 | 0.0337 | 0.0336 | 0.0337 | 0.0372 | 0.0372 | 0.0372 |
| 12 | 0.0646 | 0.0646 | 0.0646 | 0.0399 | 0.04 | 0.0399 | 0.0335 | 0.0335 | 0.0336 | 0.0371 | 0.0371 | 0.0371 |
| 13 | 0.0644 | 0.0644 | 0.0644 | 0.0398 | 0.0399 | 0.0399 | 0.0335 | 0.0334 | 0.0335 | 0.0369 | 0.037 | 0.037 |
| 14 | 0.0642 | 0.0642 | 0.0642 | 0.0397 | 0.0398 | 0.0398 | 0.0333 | 0.0333 | 0.0334 | 0.0369 | 0.0369 | 0.0369 |
| 15 | 0.064 | 0.064 | 0.064 | 0.0397 | 0.0398 | 0.0397 | 0.0333 | 0.0333 | 0.0333 | 0.0368 | 0.0368 | 0.0368 |
| 16 | 0.0639 | 0.0639 | 0.0639 | 0.0396 | 0.0397 | 0.0397 | 0.0332 | 0.0332 | 0.0332 | 0.0367 | 0.0367 | 0.0367 |
| 17 | 0.0638 | 0.0638 | 0.0638 | 0.0396 | 0.0396 | 0.0396 | 0.0331 | 0.0331 | 0.0332 | 0.0366 | 0.0366 | 0.0367 |
| 18 | 0.0637 | 0.0637 | 0.0637 | 0.0395 | 0.0396 | 0.0396 | 0.033 | 0.0331 | 0.0331 | 0.0366 | 0.0366 | 0.0366 |
| 19 | 0.0635 | 0.0636 | 0.0635 | 0.0394 | 0.0395 | 0.0395 | 0.033 | 0.033 | 0.0331 | 0.0365 | 0.0365 | 0.0365 |
| 20 | 0.0635 | 0.0635 | 0.0634 | 0.0394 | 0.0395 | 0.0395 | 0.0329 | 0.0329 | 0.033 | 0.0365 | 0.0365 | 0.0365 |
| 21 | 0.0634 | 0.0634 | 0.0633 | 0.0393 | 0.0394 | 0.0394 | 0.0329 | 0.0329 | 0.033 | 0.0364 | 0.0364 | 0.0364 |
| 22 | 0.0633 | 0.0633 | 0.0633 | 0.0393 | 0.0393 | 0.0394 | 0.0329 | 0.0329 | 0.0329 | 0.0364 | 0.0364 | 0.0364 |
| 23 | 0.0632 | 0.0631 | 0.0632 | 0.0393 | 0.0393 | 0.0393 | 0.0328 | 0.0328 | 0.0329 | 0.0363 | 0.0363 | 0.0363 |
| 24 | 0.0631 | 0.0631 | 0.0631 | 0.0392 | 0.0393 | 0.0393 | 0.0328 | 0.0328 | 0.0328 | 0.0363 | 0.0363 | 0.0363 |
| 25 | 0.063 | 0.063 | 0.063 | 0.0392 | 0.0392 | 0.0393 | 0.0327 | 0.0328 | 0.0328 | 0.0362 | 0.0362 | 0.0362 |
| 26 | 0.0629 | 0.0629 | 0.0629 | 0.0391 | 0.0392 | 0.0392 | 0.0327 | 0.0327 | 0.0328 | 0.0362 | 0.0362 | 0.0362 |
| 27 | 0.0628 | 0.0628 | 0.0628 | 0.0391 | 0.0392 | 0.0392 | 0.0327 | 0.0327 | 0.0327 | 0.0362 | 0.0362 | 0.0362 |
| 28 | 0.0628 | 0.0628 | 0.0628 | 0.0391 | 0.0391 | 0.0392 | 0.0327 | 0.0327 | 0.0327 | 0.0361 | 0.0361 | 0.0361 |
| 29 | 0.0627 | 0.0627 | 0.0627 | 0.039 | 0.0391 | 0.0392 | 0.0326 | 0.0326 | 0.0327 | 0.0361 | 0.0361 | 0.0361 |
| 30 | 0.0626 | 0.0626 | 0.0626 | 0.039 | 0.0391 | 0.0391 | 0.0326 | 0.0326 | 0.0327 | 0.036 | 0.036 | 0.0361 |
| 31 | 0.0626 | 0.0626 | 0.0626 | 0.039 | 0.0391 | 0.0391 | 0.0326 | 0.0326 | 0.0327 | 0.036 | 0.036 | 0.036 |
| 32 | 0.0626 | 0.0625 | 0.0625 | 0.039 | 0.0391 | 0.0391 | 0.0326 | 0.0326 | 0.0327 | 0.036 | 0.036 | 0.036 |
| 33 | 0.0624 | 0.0624 | 0.0624 | 0.039 | 0.039 | 0.039 | 0.0326 | 0.0326 | 0.0326 | 0.036 | 0.0359 | 0.036 |
| 34 | 0.0624 | 0.0624 | 0.0624 | 0.0389 | 0.039 | 0.039 | 0.0326 | 0.0326 | 0.0326 | 0.0359 | 0.0359 | 0.036 |
| 35 | 0.0624 | 0.0623 | 0.0624 | 0.0389 | 0.039 | 0.039 | 0.0325 | 0.0325 | 0.0326 | 0.0359 | 0.0359 | 0.0359 |
| 36 | 0.0623 | 0.0623 | 0.0623 | 0.0389 | 0.039 | 0.039 | 0.0325 | 0.0325 | 0.0348 | 0.0359 | 0.0359 | 0.0359 |
| 37 | 0.0623 | 0.0622 | 0.0622 | 0.0389 | 0.039 | 0.039 | 0.0325 | 0.0325 | 0.0339 | 0.0358 | 0.0358 | 0.0359 |
| 38 | 0.0622 | 0.0622 | 0.0622 | 0.0389 | 0.039 | 0.039 | 0.0325 | 0.0325 | 0.0337 | 0.0358 | 0.0358 | 0.0359 |
| 39 | 0.0622 | 0.0622 | 0.0621 | 0.0389 | 0.039 | 0.039 | 0.0325 | 0.0325 | 0.033 | 0.0358 | 0.0358 | 0.0358 |
| 40 | 0.0621 | 0.0621 | 0.0621 | 0.0389 | 0.039 | 0.039 | 0.0326 | 0.0325 | 0.0335 | 0.0358 | 0.0358 | 0.0358 |
| 41 | 0.0621 | 0.0621 | 0.0621 | 0.0389 | 0.039 | 0.039 | 0.0338 | 0.0325 | 0.0328 | 0.0358 | 0.0358 | 0.0358 |
| 42 | 0.0621 | 0.0621 | 0.062 | 0.0389 | 0.0389 | 0.039 | 0.0355 | 0.0325 | 0.0331 | 0.0357 | 0.0357 | 0.0358 |
| 43 | 0.0621 | 0.062 | 0.062 | 0.0389 | 0.0389 | 0.0389 | 0.0347 | 0.0325 | 0.0327 | 0.0357 | 0.0357 | 0.0358 |
| 44 | 0.062 | 0.062 | 0.062 | 0.0389 | 0.0389 | 0.0389 | 0.0355 | 0.0326 | 0.0476 | 0.0357 | 0.0357 | 0.0357 |
| 45 | 0.062 | 0.062 | 0.062 | 0.0389 | 0.0389 | 0.0389 | 0.0396 | 0.0327 | 0.034 | 0.0357 | 0.0357 | 0.0357 |
| 46 | 0.0619 | 0.062 | 0.0619 | 0.0389 | 0.0389 | 0.0389 | 0.0419 | 0.0393 | 0.04 | 0.0357 | 0.0357 | 0.0357 |
| 47 | 0.0619 | 0.0619 | 0.0619 | 0.0389 | 0.0389 | 0.0389 | 0.0369 | 0.0565 | 0.0362 | 0.0357 | 0.0357 | 0.0357 |
| 48 | 0.0619 | 0.0619 | 0.0619 | 0.0389 | 0.0389 | 0.0389 | 0.0393 | 0.0591 | 0.0428 | 0.0357 | 0.0357 | 0.0357 |
| 49 | 0.0619 | 0.0619 | 0.0619 | 0.0389 | 0.0389 | 0.0389 | 0.0376 | 0.0535 | 0.0477 | 0.0356 | 0.0357 | 0.0357 |
| 50 | 0.0619 | 0.0619 | 0.0618 | 0.0389 | 0.0389 | 0.0389 | 0.0484 | 0.0747 | 0.0535 | 0.0356 | 0.0356 | 0.0357 |
| 51 | 0.0618 | 0.0619 | 0.0618 | 0.0389 | 0.0389 | 0.0389 | 0.0644 | 0.064 | 0.0404 | 0.0356 | 0.0356 | 0.0356 |
| 52 | 0.0618 | 0.0619 | 0.0618 | 0.0389 | 0.0389 | 0.0389 | 0.0642 | 0.0521 | 0.0853 | 0.0356 | 0.0356 | 0.0356 |
| 53 | 0.0618 | 0.0618 | 0.0618 | 0.0389 | 0.0389 | 0.0389 | 0.0517 | 0.067 | 0.0893 | 0.0356 | 0.0356 | 0.0356 |
| 54 | 0.0618 | 0.0618 | 0.0618 | 0.0389 | 0.0389 | 0.0389 | 0.0517 | 0.1087 | 0.0917 | 0.0356 | 0.0356 | 0.0356 |
| 55 | 0.0618 | 0.0618 | 0.0618 | 0.0389 | 0.039 | 0.0389 | 0.0551 | 0.0947 | 0.1171 | 0.0356 | 0.0356 | 0.0356 |
| 56 | 0.0618 | 0.0618 | 0.0618 | 0.0389 | 0.039 | 0.0389 | 0.0606 | 0.0859 | 0.1342 | 0.0356 | 0.0356 | 0.0356 |
| 57 | 0.0618 | 0.0618 | 0.0617 | 0.0389 | 0.0389 | 0.0389 | 0.0702 | 0.1263 | 0.1116 | 0.0356 | 0.0356 | 0.0356 |
| 58 | 0.0618 | 0.0617 | 0.0617 | 0.0409 | 0.0389 | 0.0389 | 0.0799 | 0.1017 | 0.1212 | 0.0356 | 0.0356 | 0.0356 |
| 59 | 0.0617 | 0.0617 | 0.0617 | 0.0419 | 0.0393 | 0.0389 | 0.1121 | 0.0972 | 0.1144 | 0.0356 | 0.0356 | 0.0356 |
| 60 | 0.0617 | 0.0617 | 0.0617 | 0.0422 | 0.039 | 0.039 | 0.0943 | 0.206 | 0.1226 | 0.0356 | 0.0356 | 0.0356 |
| 61 | 0.0617 | 0.0617 | 0.0617 | 0.0446 | 0.039 | 0.039 | 0.0914 | 0.2296 | 0.168 | 0.0355 | 0.0356 | 0.0356 |
| 62 | 0.0617 | 0.0617 | 0.0617 | 0.0417 | 0.0389 | 0.0396 | 0.1466 | 0.2235 | 0.1795 | 0.0355 | 0.0355 | 0.0356 |
| 63 | 0.0652 | 0.0617 | 0.0617 | 0.0416 | 0.0618 | 0.0399 | 0.2471 | 0.2111 | 0.2042 | 0.0355 | 0.0391 | 0.0356 |
| 64 | 0.0617 | 0.0617 | 0.0617 | 0.0544 | 0.0577 | 0.0403 | 0.2359 | 0.2464 | 0.1949 | 0.0355 | 0.0371 | 0.0355 |
| 65 | 0.0617 | 0.0617 | 0.0617 | 0.0601 | 0.0403 | 0.039 | 0.2092 | 0.2239 | 0.2217 | 0.0355 | 0.039 | 0.0355 |
| 66 | 0.0617 | 0.0617 | 0.0617 | 0.06 | 0.039 | 0.039 | 0.2665 | 0.2631 | 0.2494 | 0.0355 | 0.0445 | 0.0355 |
| 67 | 0.0617 | 0.0618 | 0.0616 | 0.0618 | 0.0522 | 0.0391 | 0.2639 | 0.3127 | 0.2099 | 0.0355 | 0.0398 | 0.0355 |
| 68 | 0.0617 | 0.0619 | 0.064 | 0.0608 | 0.0399 | 0.0393 | 0.2702 | 0.3294 | 0.2526 | 0.0355 | 0.0371 | 0.0355 |
| 69 | 0.0617 | 0.0701 | 0.0625 | 0.0579 | 0.0534 | 0.0454 | 0.302 | 0.3123 | 0.2431 | 0.0355 | 0.0421 | 0.0356 |
| 70 | 0.0617 | 0.0705 | 0.0617 | 0.0675 | 0.0391 | 0.039 | 0.3537 | 0.3259 | 0.2759 | 0.0355 | 0.0404 | 0.0355 |
| 71 | 0.0638 | 0.0691 | 0.0619 | 0.0799 | 0.04 | 0.0401 | 0.3951 | 0.2852 | 0.2674 | 0.0355 | 0.0391 | 0.0355 |
| 72 | 0.0642 | 0.066 | 0.0655 | 0.0927 | 0.0446 | 0.0395 | 0.3885 | 0.2707 | 0.2844 | 0.0355 | 0.0385 | 0.0355 |
| 73 | 0.0718 | 0.0629 | 0.0672 | 0.1025 | 0.0434 | 0.0393 | 0.3651 | 0.3261 | 0.3887 | 0.0356 | 0.0385 | 0.0355 |
| 74 | 0.0702 | 0.0616 | 0.069 | 0.1223 | 0.0518 | 0.0522 | 0.398 | 0.3771 | 0.4757 | 0.043 | 0.0384 | 0.0358 |
| 75 | 0.0724 | 0.0616 | 0.0634 | 0.1249 | 0.0505 | 0.0525 | 0.4101 | 0.3591 | 0.4559 | 0.0403 | 0.0373 | 0.0357 |
| 76 | 0.073 | 0.0617 | 0.0704 | 0.1402 | 0.0577 | 0.0534 | 0.4565 | 0.372 | 0.5244 | 0.047 | 0.0368 | 0.0367 |
| 77 | 0.066 | 0.0635 | 0.0705 | 0.1568 | 0.0684 | 0.0556 | 0.5064 | 0.3957 | 0.4898 | 0.0395 | 0.0355 | 0.0355 |
| 78 | 0.0806 | 0.066 | 0.074 | 0.1739 | 0.0923 | 0.0764 | 0.5074 | 0.4629 | 0.5463 | 0.0356 | 0.0375 | 0.0355 |
| 79 | 0.0777 | 0.0696 | 0.0672 | 0.1816 | 0.0913 | 0.0897 | 0.6033 | 0.5461 | 0.5791 | 0.0356 | 0.0369 | 0.0357 |
| 80 | 0.0771 | 0.0732 | 0.07 | 0.1914 | 0.0944 | 0.0671 | 0.6633 | 0.5012 | 0.5478 | 0.0473 | 0.0366 | 0.0363 |

Table 4: **Test Mean Squared Error (MSE) by Modality and Replicate for the PBMC dataset.** Each column shows the test reconstruction error from cross-validated NMF across three random replicates per modality. Highlighted values indicate the **best (lowest) test MSE rank** for each replicate, corresponding to the optimal number of latent factors ($k^*$).

| | Added | | | Spliced Only | | | Unspliced Only | | | Concatenated | | |
|---|---|---|---|---|---|---|---|---|---|---|---|---|
| k | Rep 1 | Rep 2 | Rep 3 | Rep 1 | Rep 2 | Rep 3 | Rep 1 | Rep 2 | Rep3 | Rep 1 | Rep 2 | Rep3 |
| 2 | 0.0576 | 0.0576 | 0.0576 | 0.0242 | 0.0242 | 0.0242 | 0.0404 | 0.0404 | 0.0405 | 0.0327 | 0.0327 | 0.0327 |
| 3 | 0.0557 | 0.0557 | 0.0557 | 0.0237 | 0.0236 | 0.0236 | 0.0391 | 0.0392 | 0.0393 | 0.0318 | 0.0318 | 0.0318 |
| 4 | 0.0548 | 0.0546 | 0.0546 | 0.0232 | 0.0231 | 0.0231 | 0.0386 | 0.0387 | 0.0388 | 0.0313 | 0.0313 | 0.0312 |
| 5 | 0.0539 | 0.0539 | 0.0539 | 0.0228 | 0.0228 | 0.0229 | 0.0384 | 0.0385 | 0.0386 | 0.0308 | 0.0309 | 0.0309 |
| 6 | 0.0536 | 0.0535 | 0.0536 | 0.0227 | 0.0226 | 0.0226 | 0.0383 | 0.0383 | 0.0384 | 0.0307 | 0.0307 | 0.0307 |
| 7 | 0.0531 | 0.0531 | 0.0531 | 0.0225 | 0.0225 | 0.0225 | 0.0381 | 0.0382 | 0.0383 | 0.0306 | 0.0305 | 0.0305 |
| 8 | 0.0529 | 0.0529 | 0.0528 | 0.0224 | 0.0224 | 0.0224 | 0.038 | 0.0381 | 0.0382 | 0.0303 | 0.0304 | 0.0304 |
| 9 | 0.0527 | 0.0526 | 0.0526 | 0.0223 | 0.0223 | 0.0223 | 0.0379 | 0.038 | 0.0381 | 0.0302 | 0.0303 | 0.0302 |
| 10 | 0.0525 | 0.0525 | 0.0525 | 0.0223 | 0.0223 | 0.0223 | 0.0379 | 0.0379 | 0.038 | 0.0301 | 0.0302 | 0.0302 |
| 11 | 0.0524 | 0.0524 | 0.0524 | 0.0223 | 0.0222 | 0.0222 | 0.0378 | 0.0379 | 0.038 | 0.0301 | 0.0301 | 0.0301 |
| 12 | 0.0523 | 0.0523 | 0.0523 | 0.0222 | 0.0222 | 0.0222 | 0.0377 | 0.0378 | 0.0379 | 0.0301 | 0.0301 | 0.0301 |
| 13 | 0.0522 | 0.0522 | 0.0522 | 0.0222 | 0.0221 | 0.0221 | 0.0377 | 0.0378 | 0.0379 | 0.03 | 0.0301 | 0.03 |
| 14 | 0.0522 | 0.0521 | 0.0521 | 0.0221 | 0.0221 | 0.0221 | 0.0377 | 0.0378 | 0.0379 | 0.03 | 0.03 | 0.03 |
| 15 | 0.0521 | 0.0521 | 0.0521 | 0.0221 | 0.0221 | 0.0221 | 0.0377 | 0.0377 | 0.0378 | 0.0299 | 0.03 | 0.0299 |
| 16 | 0.052 | 0.052 | 0.052 | 0.0221 | 0.0221 | 0.022 | 0.0376 | 0.0377 | 0.0378 | 0.0299 | 0.0299 | 0.0299 |
| 17 | 0.052 | 0.052 | 0.052 | 0.0221 | 0.0221 | 0.022 | 0.0376 | 0.0377 | 0.0377 | 0.0299 | 0.0299 | 0.0299 |
| 18 | 0.0519 | 0.0519 | 0.0519 | 0.0221 | 0.0221 | 0.022 | 0.0376 | 0.0377 | 0.0377 | 0.0298 | 0.0299 | 0.0299 |
| 19 | 0.0519 | 0.0518 | 0.0519 | 0.0221 | 0.022 | 0.022 | 0.0376 | 0.0377 | 0.0377 | 0.0298 | 0.0299 | 0.0298 |
| 20 | 0.0518 | 0.0518 | 0.0518 | 0.022 | 0.022 | 0.022 | 0.0375 | 0.0377 | 0.0377 | 0.0298 | 0.0299 | 0.0298 |
| 21 | 0.0518 | 0.0518 | 0.0518 | 0.022 | 0.022 | 0.022 | 0.0375 | 0.0376 | 0.0377 | 0.0298 | 0.0298 | 0.0298 |
| 22 | 0.0518 | 0.0518 | 0.0518 | 0.022 | 0.022 | 0.022 | 0.0376 | 0.0376 | 0.0377 | 0.0298 | 0.0298 | 0.0298 |
| 23 | 0.0518 | 0.0517 | 0.0518 | 0.022 | 0.022 | 0.022 | 0.0376 | 0.0376 | 0.0377 | 0.0298 | 0.0298 | 0.0298 |
| 24 | 0.0517 | 0.0517 | 0.0517 | 0.022 | 0.022 | 0.022 | 0.0375 | 0.0377 | 0.0377 | 0.0298 | 0.0298 | 0.0298 |
| 25 | 0.0517 | 0.0517 | 0.0517 | 0.022 | 0.022 | 0.022 | 0.0375 | 0.0377 | 0.0376 | 0.0297 | 0.0298 | 0.0297 |
| 26 | 0.0517 | 0.0517 | 0.0517 | 0.0264 | 0.022 | 0.022 | 0.0375 | 0.0377 | 0.0416 | 0.0297 | 0.0298 | 0.0297 |
| 27 | 0.0517 | 0.0516 | 0.0516 | 0.022 | 0.022 | 0.0219 | 0.0395 | 0.0382 | 0.0454 | 0.0297 | 0.0298 | 0.0297 |
| 28 | 0.0517 | 0.0516 | 0.0516 | 0.022 | 0.022 | 0.0219 | 0.0489 | 0.0379 | 0.0494 | 0.0297 | 0.0298 | 0.0297 |
| 29 | 0.0516 | 0.0516 | 0.0516 | 0.022 | 0.022 | 0.0253 | 0.0493 | 0.0382 | 0.0547 | 0.0297 | 0.0297 | 0.0297 |
| 30 | 0.0517 | 0.0516 | 0.0516 | 0.022 | 0.022 | 0.0221 | 0.045 | 0.0398 | 0.0565 | 0.0297 | 0.0297 | 0.0297 |
| 31 | 0.0516 | 0.0516 | 0.0516 | 0.022 | 0.022 | 0.0292 | 0.0377 | 0.0411 | 0.06 | 0.0297 | 0.0298 | 0.0297 |
| 32 | 0.0516 | 0.0516 | 0.0517 | 0.022 | 0.0222 | 0.024 | 0.0451 | 0.0401 | 0.0484 | 0.0297 | 0.0297 | 0.0297 |
| 33 | 0.0516 | 0.0515 | 0.0517 | 0.0221 | 0.0221 | 0.037 | 0.0386 | 0.0386 | 0.0484 | 0.0297 | 0.033 | 0.0297 |
| 34 | 0.0516 | 0.0515 | 0.0516 | 0.0222 | 0.0221 | 0.0384 | 0.0441 | 0.0395 | 0.0615 | 0.0297 | 0.0357 | 0.0297 |
| 35 | 0.0516 | 0.0515 | 0.0516 | 0.0222 | 0.0221 | 0.0418 | 0.0437 | 0.0549 | 0.0621 | 0.0297 | 0.0357 | 0.0297 |
| 36 | 0.0516 | 0.0515 | 0.0515 | 0.0222 | 0.0223 | 0.0375 | 0.0446 | 0.0498 | 0.0685 | 0.033 | 0.039 | 0.0297 |
| 37 | 0.0515 | 0.0515 | 0.0515 | 0.0222 | 0.0223 | 0.0378 | 0.0487 | 0.0698 | 0.0728 | 0.0367 | 0.0373 | 0.0297 |
| 38 | 0.0516 | 0.0515 | 0.0515 | 0.0224 | 0.0223 | 0.0401 | 0.0551 | 0.0818 | 0.0761 | 0.037 | 0.0387 | 0.0297 |
| 39 | 0.0515 | 0.0515 | 0.0515 | 0.0224 | 0.0223 | 0.0465 | 0.0607 | 0.0787 | 0.0741 | 0.0365 | 0.0382 | 0.0305 |
| 40 | 0.0515 | 0.0515 | 0.0516 | 0.0224 | 0.0228 | 0.0504 | 0.0622 | 0.0788 | 0.0783 | 0.0297 | 0.0381 | 0.0302 |
| 41 | 0.0521 | 0.0515 | 0.0515 | 0.0412 | 0.0227 | 0.0452 | 0.0463 | 0.0831 | 0.0856 | 0.0367 | 0.036 | 0.0298 |
| 42 | 0.0549 | 0.0516 | 0.0516 | 0.0588 | 0.0227 | 0.0361 | 0.0523 | 0.0915 | 0.0803 | 0.0348 | 0.0363 | 0.0298 |
| 43 | 0.0525 | 0.0516 | 0.0516 | 0.0611 | 0.0228 | 0.0319 | 0.083 | 0.0879 | 0.0878 | 0.0434 | 0.037 | 0.0298 |
| 44 | 0.0537 | 0.0523 | 0.0516 | 0.0571 | 0.0226 | 0.0312 | 0.0943 | 0.1018 | 0.0942 | 0.0377 | 0.036 | 0.0298 |
| 45 | 0.0551 | 0.0533 | 0.0516 | 0.0539 | 0.0223 | 0.0342 | 0.0896 | 0.112 | 0.0921 | 0.0318 | 0.0384 | 0.0297 |
| 46 | 0.0544 | 0.0537 | 0.0619 | 0.0729 | 0.0229 | 0.0512 | 0.0848 | 0.1154 | 0.0912 | 0.0389 | 0.035 | 0.0298 |
| 47 | 0.0561 | 0.0537 | 0.0657 | 0.0458 | 0.0235 | 0.0382 | 0.0859 | 0.1265 | 0.1134 | 0.032 | 0.0298 | 0.0298 |
| 48 | 0.0588 | 0.0541 | 0.0646 | 0.0498 | 0.0234 | 0.0413 | 0.1177 | 0.1325 | 0.1207 | 0.0301 | 0.0352 | 0.0304 |
| 49 | 0.0572 | 0.0559 | 0.0681 | 0.0441 | 0.024 | 0.0454 | 0.1117 | 0.1378 | 0.1246 | 0.0302 | 0.0398 | 0.0306 |
| 50 | 0.0568 | 0.0535 | 0.0724 | 0.0301 | 0.0238 | 0.0436 | 0.1052 | 0.1506 | 0.11 | 0.0308 | 0.0422 | 0.0304 |
| 51 | 0.0644 | 0.0537 | 0.0761 | 0.05 | 0.0245 | 0.0439 | 0.1204 | 0.1349 | 0.1208 | 0.0311 | 0.0431 | 0.0336 |
| 52 | 0.0655 | 0.0541 | 0.0664 | 0.0495 | 0.0246 | 0.0447 | 0.1269 | 0.1418 | 0.1302 | 0.0314 | 0.0483 | 0.0305 |
| 53 | 0.0762 | 0.0566 | 0.0731 | 0.0575 | 0.026 | 0.0561 | 0.1145 | 0.1558 | 0.1359 | 0.031 | 0.0456 | 0.0307 |
| 54 | 0.0982 | 0.0563 | 0.0822 | 0.0594 | 0.0243 | 0.0895 | 0.1117 | 0.16 | 0.1447 | 0.0313 | 0.052 | 0.0307 |
| 55 | 0.1014 | 0.0571 | 0.0825 | 0.1701 | 0.0252 | 0.1324 | 0.1281 | 0.1401 | 0.1664 | 0.0318 | 0.0583 | 0.0307 |
| 56 | 0.1001 | 0.0567 | 0.0836 | 0.133 | 0.0277 | 0.1458 | 0.1875 | 0.1761 | 0.1637 | 0.0334 | 0.0663 | 0.0307 |
| 57 | 0.1032 | 0.0689 | 0.0909 | 0.1088 | 0.0234 | 0.157 | 0.1929 | 0.1711 | 0.1897 | 0.033 | 0.0686 | 0.0308 |
| 58 | 0.1144 | 0.075 | 0.0961 | 0.0972 | 0.0251 | 0.1668 | 0.195 | 0.1821 | 0.1843 | 0.0335 | 0.071 | 0.031 |
| 59 | 0.1313 | 0.0668 | 0.0972 | 0.0944 | 0.0254 | 0.1585 | 0.2145 | 0.1828 | 0.1948 | 0.0345 | 0.0694 | 0.0316 |
| 60 | 0.1333 | 0.0999 | 0.0995 | 0.0998 | 0.0565 | 0.1452 | 0.2056 | 0.1921 | 0.2902 | 0.0349 | 0.0755 | 0.0315 |
| 61 | 0.106 | 0.0697 | 0.1034 | 0.1131 | 0.101 | 0.1632 | 0.2369 | 0.2184 | 0.3503 | 0.0352 | 0.074 | 0.0319 |
| 62 | 0.1369 | 0.0717 | 0.104 | 0.1011 | 0.1043 | 0.1781 | 0.1923 | 0.2764 | 0.3681 | 0.0404 | 0.0571 | 0.0347 |
| 63 | 0.1571 | 0.0736 | 0.1124 | 0.099 | 0.1152 | 0.1978 | 0.1921 | 0.2798 | 0.3844 | 0.0387 | 0.0882 | 0.0463 |
| 64 | 0.1662 | 0.074 | 0.1142 | 0.168 | 0.1257 | 0.2039 | 0.2204 | 0.3073 | 0.402 | 0.0436 | 0.0821 | 0.0379 |
| 65 | 0.109 | 0.0662 | 0.1132 | 0.1743 | 0.1221 | 0.2114 | 0.2616 | 0.3012 | 0.4472 | 0.0401 | 0.0928 | 0.044 |
| 66 | 0.1065 | 0.108 | 0.1075 | 0.19 | 0.0627 | 0.2084 | 0.2449 | 0.2504 | 0.3901 | 0.0423 | 0.0907 | 0.0527 |
| 67 | 0.1177 | 0.1017 | 0.1077 | 0.2131 | 0.1604 | 0.2254 | 0.2761 | 0.2852 | 0.4327 | 0.0458 | 0.0872 | 0.0541 |
| 68 | 0.1183 | 0.1029 | 0.1172 | 0.2092 | 0.1751 | 0.2467 | 0.3223 | 0.3353 | 0.4173 | 0.0445 | 0.0898 | 0.045 |
| 69 | 0.1336 | 0.1178 | 0.1158 | 0.2217 | 0.1845 | 0.2857 | 0.3218 | 0.3396 | 0.4241 | 0.0482 | 0.0713 | 0.0437 |
| 70 | 0.1509 | 0.1176 | 0.1126 | 0.2208 | 0.2005 | 0.2887 | 0.3457 | 0.3525 | 0.4541 | 0.0558 | 0.0707 | 0.0669 |
| 71 | 0.1642 | 0.1523 | 0.0847 | 0.2257 | 0.2082 | 0.2947 | 0.3673 | 0.3752 | 0.4572 | 0.0554 | 0.0905 | 0.0714 |
| 72 | 0.1898 | 0.1534 | 0.0955 | 0.2332 | 0.2087 | 0.3091 | 0.3383 | 0.309 | 0.4765 | 0.056 | 0.0965 | 0.1002 |
| 73 | 0.217 | 0.1614 | 0.1008 | 0.26 | 0.1659 | 0.3086 | 0.3415 | 0.3428 | 0.4786 | 0.0575 | 0.1161 | 0.101 |
| 74 | 0.1193 | 0.1577 | 0.1203 | 0.2623 | 0.1673 | 0.323 | 0.4292 | 0.3768 | 0.4935 | 0.0532 | 0.1076 | 0.0751 |
| 75 | 0.1999 | 0.0958 | 0.1015 | 0.2543 | 0.209 | 0.3429 | 0.4482 | 0.3942 | 0.4619 | 0.0678 | 0.109 | 0.0981 |
| 76 | 0.2257 | 0.0926 | 0.1377 | 0.287 | 0.2467 | 0.3716 | 0.4802 | 0.3997 | 0.4404 | 0.0908 | 0.1401 | 0.0936 |
| 77 | 0.155 | 0.0912 | 0.1872 | 0.3049 | 0.2551 | 0.3677 | 0.5158 | 0.431 | 0.5195 | 0.0922 | 0.1295 | 0.0945 |
| 78 | 0.2303 | 0.105 | 0.1923 | 0.3174 | 0.2552 | 0.3846 | 0.511 | 0.4026 | 0.5764 | 0.1018 | 0.1234 | 0.1044 |
| 79 | 0.2269 | 0.1177 | 0.1964 | 0.3214 | 0.2486 | 0.4172 | 0.5296 | 0.4758 | 0.5369 | 0.1099 | 0.1194 | 0.1098 |
| 80 | 0.3064 | 0.1378 | 0.1187 | 0.3327 | 0.2713 | 0.4977 | 0.5478 | 0.4579 | 0.6097 | 0.1146 | 0.1271 | 0.1129 |
